# Haplotype-Resolved DNA Methylation at the *APOE* Locus identifies Allele-Specific Epigenetic Signatures Relevant to Alzheimer’s Disease Risk

**DOI:** 10.1101/2025.07.01.662592

**Authors:** Rylee M. Genner, Melissa Meredith, Abraham Moller, Cory Weller, Kensuke Daida, Alexis Ayuketah, Pilar Alvarez Jerez, Stuart Akeson, Laksh Malik, Breeana Baker, Cedric Kouam, Kimberly Paquette, Stefano Marenco, Pavan Auluck, Ajeet Mandal, Benedict Paten, Xylena Reed, Miten Jain, Mark R. Cookson, Andrew B. Singleton, Mike Nalls, Cornelis Blauwendraat, Kimberley J. Billingsley

## Abstract

The *APOE* gene encodes a key lipid transport protein and plays a central role in Alzheimer’s disease (AD) pathogenesis. Three common *APOE* alleles, ε2 (rs7412(C>T), ε3 (reference), and ε4 (rs429358(T>C)), arise from two coding variants in exon 4 and confer distinct AD risk profiles, with ε4 increasing risk and ε2 providing protection. The ε3-linked *APOE* variant rs769455[T] has also been associated with elevated AD risk in individuals of African ancestry carrying both rs769455[T] and ε4 alleles. These single nucleotide variants (SNVs) reside in a cytosine-phosphate-guanine (CpG) island, which is a region with a higher frequency of CpG sites compared to the rest of the genome. CpG sites are subject to 5-methylcytosine (5mC) methylation by DNA methyltransferases which add a methyl group to the fifth carbon on the cytosine residue of a CpG site. The presence of SNVs can disrupt this process, making these regions prime targets for differential methylation; however, allele-specific methylation patterns in *APOE* remain poorly resolved due to technical limitations of conventional bisulfite and methylation array based methods, including degraded DNA quality, sparse CpG coverage, and lack of haplotype phasing. Here, we leverage high-accuracy long-read sequencing data to generate haplotype-resolved methylation profiles of the *APOE* locus in 332 postmortem brain samples from two ancestrally different cohorts. This includes 201 individuals of European ancestry from the North American Brain Expression Consortium (NABEC), comprising 402 haplotypes (48 ε2 and 58 ε4 alleles), and 131 individuals of African and African admixed ancestry from the Human Brain Core Collection (HBCC), comprising 262 haplotypes (25 ε2, 64 ε4, and 7 rs769455 alleles). A linear regression analysis identified 18 novel differentially methylated CpG sites (DMCs) associated with *APOE* ε2, ε4, and rs769455 within a gene cluster spanning *TOMM40, APOE, APOC1,* and *APOC4-APOC2*. This represents the most comprehensive haplotype-resolved methylation study of *APOE* in human brain tissue to date. Our results uncover distinct allele-specific methylation signatures and demonstrate the power of long-read sequencing for resolving epigenetic variation relevant to AD risk.

## Introduction

The Apolipoprotein E (*APOE*) gene, located on chromosome 19, encodes a lipid transport protein essential for neuronal development, maintenance, and repair ^1–3^. *APOE* plays a critical role in the central nervous and cardiovascular systems and is a major genetic determinant of Alzheimer’s disease (AD) risk ^4,5^. Two single nucleotide variants (SNVs) in exon 4 define the ε2 (rs7412(C>T)), ε3 (reference), and ε4 (rs429358(T>C)) alleles that modulate AD susceptibility. The ε4 allele is the strongest common genetic risk factor for late-onset AD (LOAD), increasing disease risk in a dose-dependent manner estimated to be 2-to 12-fold compared to the reference ε3 allele ^5–8^, while the ε2 allele confers a protective effect ^9^. Recently, an *APOE*-ε3-linked missense variant rs769455(C>T) was identified as a potential population-specific risk factor for AD in individuals of African ancestry who also carry the ε4 allele ^10,11^. The roles that epigenetic modifications play in disease development cannot be fully understood without considering the population specificities of methylation, and the discovery of ancestry-associated variants like rs769455[T] underscores the need for studying *APOE* genetic and epigenetic variation across genetic ancestries.

Epigenetic modifications like DNA methylation can also influence *APOE* expression and have been implicated in AD pathogenesis ^12,13^. 5-methylcytosine (5mC) refers to the transient addition of a methyl group to the fifth carbon position of a cytosine residue. This occurs most often at cytosines that are immediately adjacent to guanines, commonly called cytosine-phosphate-guanine (CpG) sites, which can be condensed within regulatory regions called CpG islands ^14–16^. The *APOE* gene cluster region, which was first defined by Cervantes et al. (2011) and spans the *TOMM40, APOE, APOC1,* and *APOC4-APOC2* genes^17–21^, contains CpG islands and other regulatory regions that may be modulating neurodegeneration by regulating gene expression and lipid metabolism in the brain^21^. Previous studies using short-read sequencing (SRS) and bisulfite sequencing have reported differentially methylated CpG sites associated with the ε2 and ε4 alleles in this cluster region ^13,22–25^. However, these approaches face several limitations: (i) bisulfite conversion degrades DNA ^26–28^; (ii) array-based methods, such as the Illumina Infinium HumanMethylation450 BeadChip array, provide incomplete CpG site coverage ^29^; and (iii) SRS generates fragmented reads that hinder accurate genome assembly and phasing ^30–32^. As a result, prior studies relied on genotype-based methylation assessments without resolving allele-specific epigenetic differences. For an overview of previous publications looking into *APOE* methylation differences in human blood and brain tissue samples, see Supplementary Table S1. Long-read sequencing (LRS) can overcome these limitations by directly detecting genome-wide nucleotide modifications at single-nucleotide resolution without requiring chemical conversion. Oxford Nanopore Technology (ONT) LRS, in particular, enables phased haplotype-resolved methylation profiling by sequencing long DNA fragments (10 kb – 1 Mb), improving genome contiguity, and enabling direct methylation detection ^33–35^. Unlike conventional approaches, LRS provides a more comprehensive view of methylation heterogeneity across the genome, making it a better method for studying allele-specific epigenetic contributions to neurodegenerative disease.

Here, we leveraged publicly available LRS data from 332 postmortem frontal cortex samples generated by the NIH Center for Alzheimer’s and Related Dementias (CARD) Long-Read Sequencing Initiative (https://card.nih.gov/research-programs/long-read-sequencing), including individuals of European (EUR) and African and African admixed (AFR) ancestry. We examined allele-specific DNA methylation associated with three *APOE* variants (ε2 and ε4 (EUR and AFR cohorts), and rs769455[T] (AFR cohort only)), across a 47,587 bp region encompassing the full *APOE* gene cluster. By resolving allele-specific methylation patterns in an ancestrally diverse dataset, our study advances the understanding of the epigenetic mechanisms linking *APOE* variation to AD risk alleles. These findings establish a framework for future functional studies aimed at elucidating how allele-specific methylation influences gene regulation in the brain.

## Results

### Comprehensive resolution of haplotypes and allele-specific methylation through long-read sequencing

We leveraged publicly available datasets from the CARD Long-read Sequencing Initiative, comprising WGS ONT sequencing data from 332 postmortem brain samples from individuals with no known neurological diseases across the NABEC (n = 201) and HBCC (n = 131) cohorts ^36^. The average age of the samples used in the analysis was 52.37 and 45.19 years old for NABEC and HBCC, respectively. 35.6% of the NABEC samples and 38.9% of the HBCC samples were from female donors. Ancestry analysis confirmed that the NABEC samples were all of European ancestry (EUR) and all HBCC samples used in the analysis were all of African and African admixed ancestry (AFR) (Supplementary Table S2). Three AD-associated SNVs located in *APOE* exon 4 (ε2, ε4, and rs769455[T]) with MAFs > 0.01 were selected for the allele-specific methylation analysis (Table 1). Allele-specific differential methylation profiling was performed at the *APOE* cluster region (chr19:44889556–44953378, hg38, identified by Cervantes et al. (2011)), which encompasses the *TOMM40*, *APOE*, *APOC1*, and *APOC4-APOC2* genes (Figure 1B). After phasing, the mean haplotype-specific coverage across this region was 12× (read N50 = 28.1 kb) in NABEC and 10.6× (read N50 = 25.6 kb) in HBCC samples (Supplementary Table S3). As expected, haplotype-level coverage was approximately halved relative to total coverage. A schematic of the pipeline used to generate haplotype-resolved variant calls and CpG methylation profiles^34^ along with representative graphs illustrating phased and unphased methylation frequency output is shown in Figure 1A.

**Figure 1.**
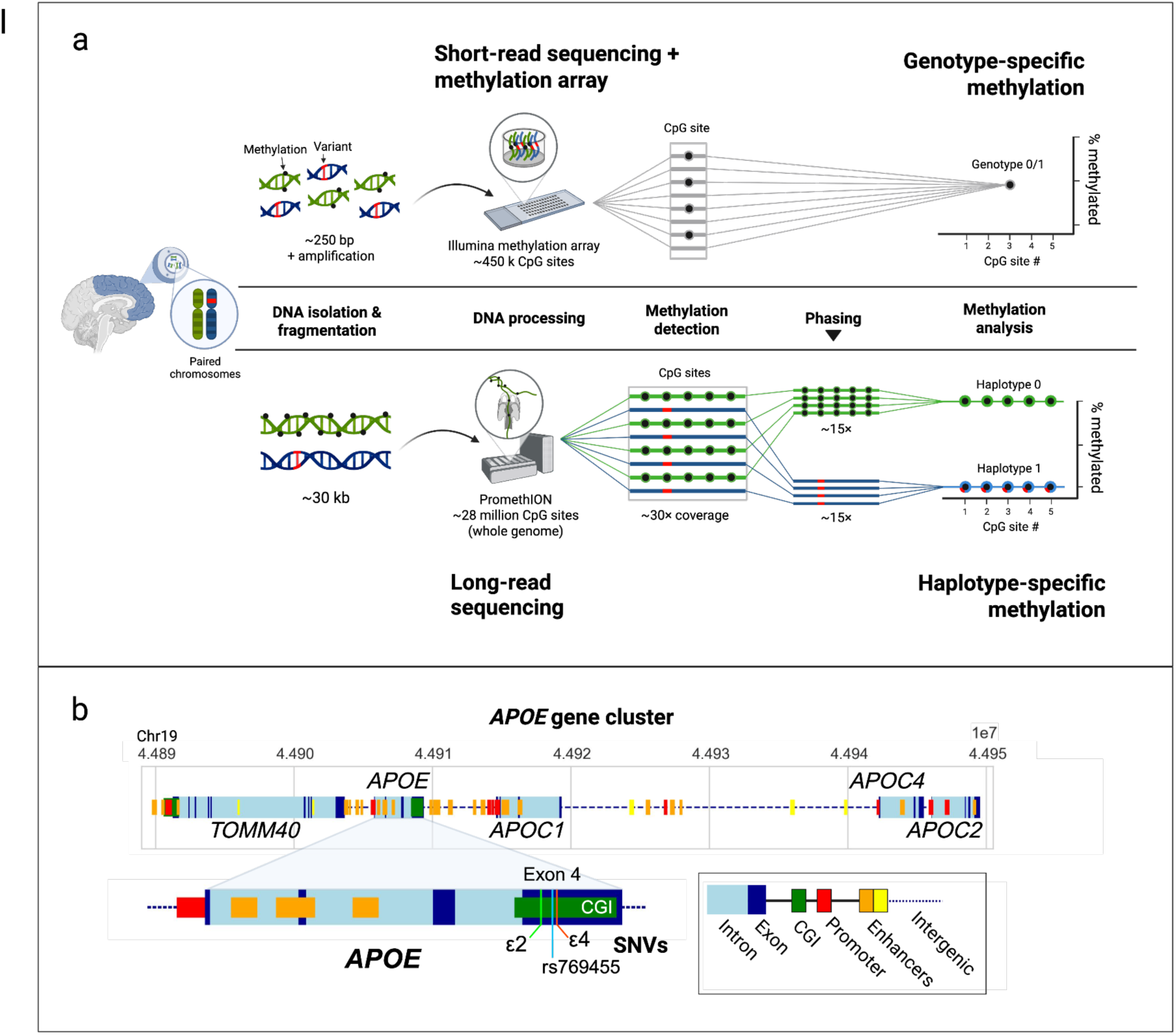
Comparison of methylation detection methods and schematic of APOE cluster region. **a)** Overview of the Illumina methylation array-based pipeline used for genotype-level methylation detection (top) and long-read sequencing pipeline allele-specific methylation analysis (bottom). **b)** Schematic of the *APOE* gene cluster, highlighting the *APOE* gene and the locations of the ε2 (rs7412), ε4 (rs429358), and rs769455[T] single nucleotide variants (SNVs) within the *APOE* exon 4 CpG island.

**Table 1.**
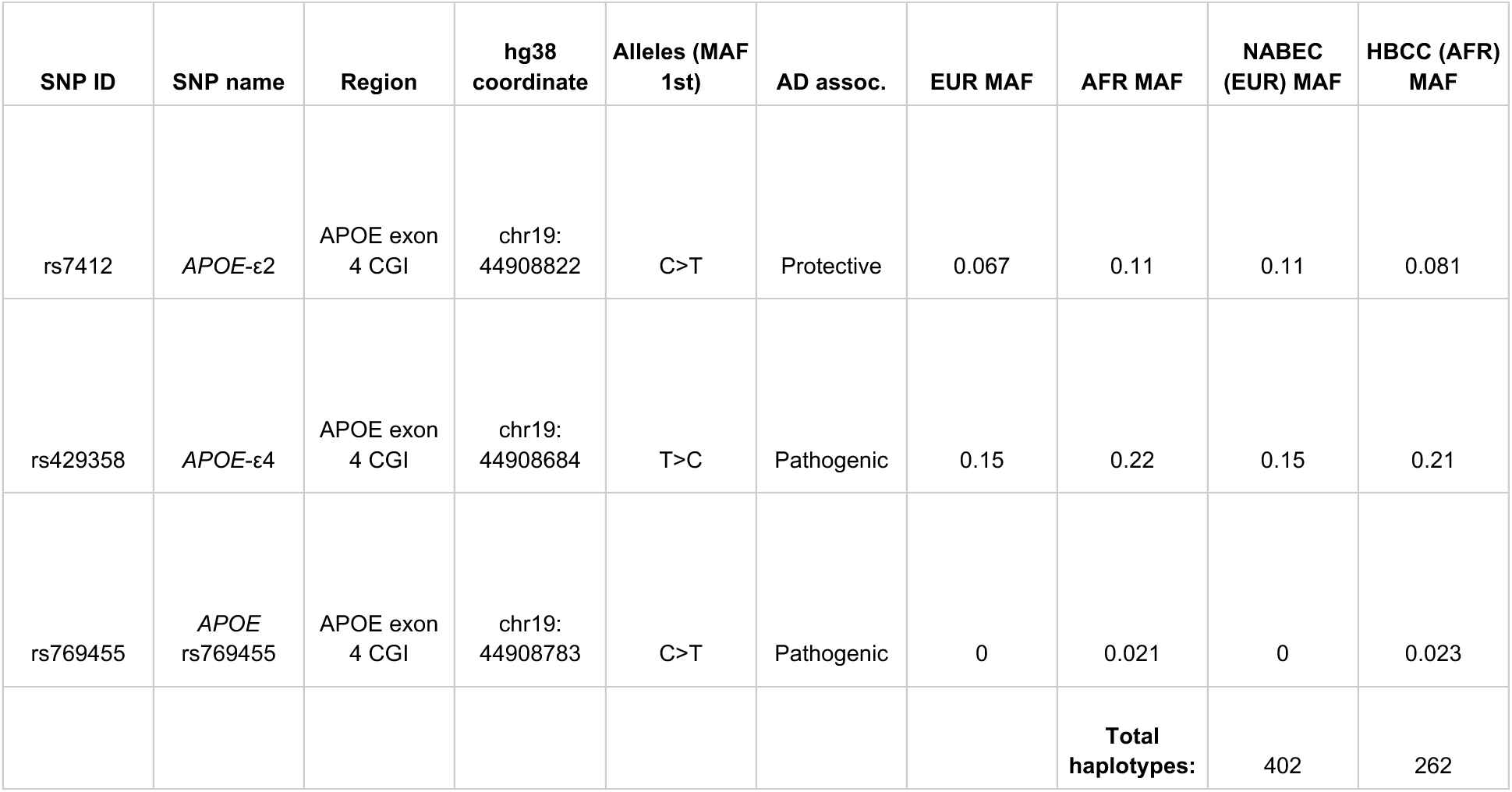
Counts and minor allele frequencies (MAFs) of the three *APOE* alleles included in the analysis. EUR (European) and AFR (African and African Admixed) MAFs were taken from Gnomad v4.1.0 using the following sample totals: ε2, ε4 and rs769455[T] European (EUR) totals = 89,919, 17,4295, and 74 respectively and ε2, ε4 and rs769455[T] African and African American (AFR) sample totals = 7,729, 16,245, and 1,549 respectively. Allele counts from the NABEC and HBCC cohorts were used to calculate variant frequencies.

| SNP ID | SNP name | Region | hg38 coordinate | Alleles (MAF 1st) | AD assoc. | EUR MAF | AFR MAF | NABEC (EUR) MAF | HBCC (AFR) MAF |
| --- | --- | --- | --- | --- | --- | --- | --- | --- | --- |
| rs7412 | *APOE*-ε2 | *APOE* exon 4 CGI | chr19: 44908822 | C>T | Protective | 0.067 | 0.11 | 0.11 | 0.081 |
| rs429358 | *APOE*-ε4 | *APOE* exon 4 CGI | chr19: 44908684 | T>C | Pathogenic | 0.15 | 0.22 | 0.15 | 0.21 |
| rs769455 | *APOE* rs769455 | *APOE* exon 4 CGI | chr19: 44908783 | C>T | Pathogenic | 0 | 0.021 | 0 | 0.023 |
|  |  |  |  |  |  |  | **Total haplotypes:** | 402 | 262 |

To evaluate the comprehensiveness of long-read methylation profiling, we compared CpG detection in our ONT data to an Illumina Infinium HumanMethylation450 BeadChip array of bulk brain tissue from the Religious Orders Study/Memory and Aging Project (ROSMAP) study, a large, well-characterized cohort of aged individuals with available postmortem brain methylation data ^37^. Long-read sequencing enabled higher-resolution methylation profiling of the *APOE* locus, substantially increasing the number of CpG sites detected compared to bisulfite sequencing. Across the *APOE* cluster region, ONT WGS data from NABEC samples (n = 201, EUR ancestry) identified 1,556 CpG sites, whereas the Illumina Infinium HumanMethylation450 BeadChip array data from ROSMAP samples (n = 697, EUR ancestry) detected 46 CpG sites (Figure 2A–B). One of the updated versions of this array is the Illumina Infinium HumanMethylation850 BeadChip array which measures methylation of over 850k CpG sites across the human genome. This array would have detected 76 CpG sites within the *APOE* gene cluster region, and these sites can be seen as blue and red lines along the “Illumina 850k” UCSC Genome track of Figure 2c. We did not have access to sequencing data from frontal cortex brain tissue samples that had been sequenced with this upgraded array, which is why the Illumina Infinium HumanMethylation450 BeadChip array data was used in this comparison. The 46 CpG sites that were detected in both European brain tissue cohorts by both platforms (Figure 2B, red points) showed highly correlated methylation frequencies (Supplementary Figure 1).

**Figure 2.**
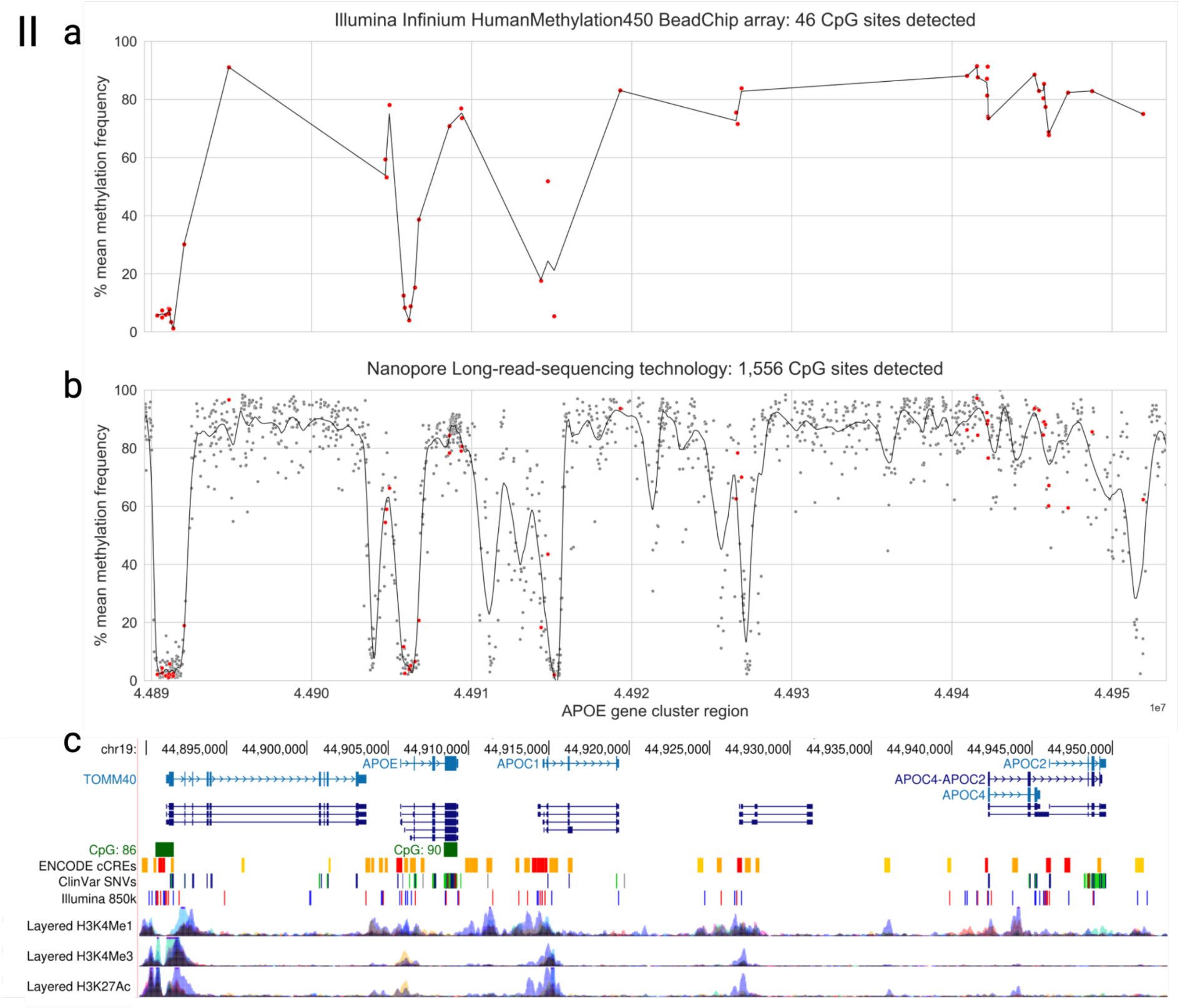
Comparison of CpG sites detected in the *APOE* gene cluster region by the Illumina Infinium HumanMethylation450 BeadChip array and ONT’s long-read sequencing technology. Plots depict the mean methylation frequencies of CpG sites across samples detected in the *APOE* cluster region (*TOMM40, APOE, APOC1, APOC4*, and *APOC2*; hg38 chr19:44889556-44953378). **a)** Mean methylation of 46 CpG sites in 697 samples from the Religious Orders Study/Memory and Aging Project, detected by the Illumina Infinium HumanMethylation450 BeadChip array. These samples came from a mix of AD and control individuals of European ancestry. **b)** Mean methylation of 1,556 CpG sites in 205 samples from the North American Brain Expression Cohort detected by ONT’s PromethION long-read sequencing data as part of the NIH CARD Long Read Initiative ^36^. CpG sites shared with the Illumina Infinium HumanMethylation450 BeadChip array are highlighted in red. **c)** UCSC Genome Browser tracks of the *APOE* cluster region, displaying GENCODE V47 genes, RefSeq transcripts, CpG islands, ENCODE Candidate Cis-Regulatory Elements, ClinVar SNVs, Illumina Infinium HumanMethylation850 BeadChip array probes, and ENCODE histone marks (H3K4Me1, H3K4Me3, H3K27Ac). Tracks align with Figure 2a and b coordinates.

CpG sites detected by bisulfite sequencing were largely restricted to promoter and exonic regions, whereas long-read profiling extended coverage to intronic, intergenic, and regulatory regions (Figure 2C) to capture a broader methylation landscape. Overall, ONT long-read sequencing detected approximately 33× more CpG sites than the Illumina Infinium HumanMethylation450 BeadChip array, underscoring the advantage of this approach for resolving previously inaccessible regions of the brain methylome.

### Haplotype-resolved long-read sequencing refines phylogenetic structure

To evaluate the contribution of haplotype phasing to the resolution of genetic structure at the *APOE* locus, we constructed dendrograms based on both phased and unphased genomic sequences and annotated them according to their *APOE* genotypes and haplotypes. All trees were rooted using the chimpanzee *APOE*-ε4 sequence, which represents the ancestral allele ^38^. The unphased dendrogram demonstrated genotype-level separation but showed discontinuous and partially overlapping clustering of alleles (Supplementary Figure 2A). In contrast, the haplotype-resolved data revealed distinct and well-separated phylogenetic clades corresponding to the *APOE* ε2, ε3, and ε4 alleles, as well as a distinct cluster for alleles harboring the rs769455[T] variant (Supplementary Figure 2B). Across the NABEC and HBCC datasets, a total of 664 phased haplotypes were resolved, enabling allele-specific analyses of methylation and sequence diversity.

Given that NABEC and HBCC samples were sequenced using different ONT chemistries (R9 and R10, respectively), we included one sample from the NABEC cohort that was sequenced using both chemistries ^35^ to confirm that clustering was driven by sequence differences rather than chemistry-specific artifacts. As expected, the *APOE* alleles in this sample clustered together, supporting the robustness of the observed phylogenetic separation (Supplementary Figure 3). These findings underscore the utility of LRS for resolving complex haplotype structure and enabling allele-specific epigenetic analyses at functionally important loci.

### Allele-specific differentially methylated CpG sites in the *APOE* gene cluster region

We identified a total of 19 allele-specific DMCs in the *APOE* gene cluster region, including 18 novel DMCs and one previously reported DMC using phased methylation data from NABEC and HBCC cohorts. In the NABEC cohort (n=201, EUR ancestry), eight CpG sites showed significant allele-specific methylation differences (BH-FDR-corrected p value < 0.05). Three sites were associated with the *ε2* allele (cpg_chr19_44914329, cpg_chr19_44914361 and cpg_chr19_44921919), and four sites were associated with the *ε4* allele (cpg_chr19_44901266, cpg_chr19_44914329, cpg_chr19_44917921, cpg_chr19_44917959, cpg chr19_44917997 and cpg_chr19_44918064). These sites were distributed across functionally relevant regions: two were located within a promoter region of the *APOC1* gene (cpg_chr19_44914329 and cpg_chr19_44914361), four overlapped a short interspersed nuclear element (SINE) within intron 3 of the *APOC1* gene (cpg_chr19_4491721, cpg_chr19_44917595, cpg_chr19_44917997 and cpg_chr19_44918064), and one was located in the intergenic region between *APOC1* and *APOE* (cpg_chr19_44921919) (Table 2; Figure 3A).

**Figure 3.**
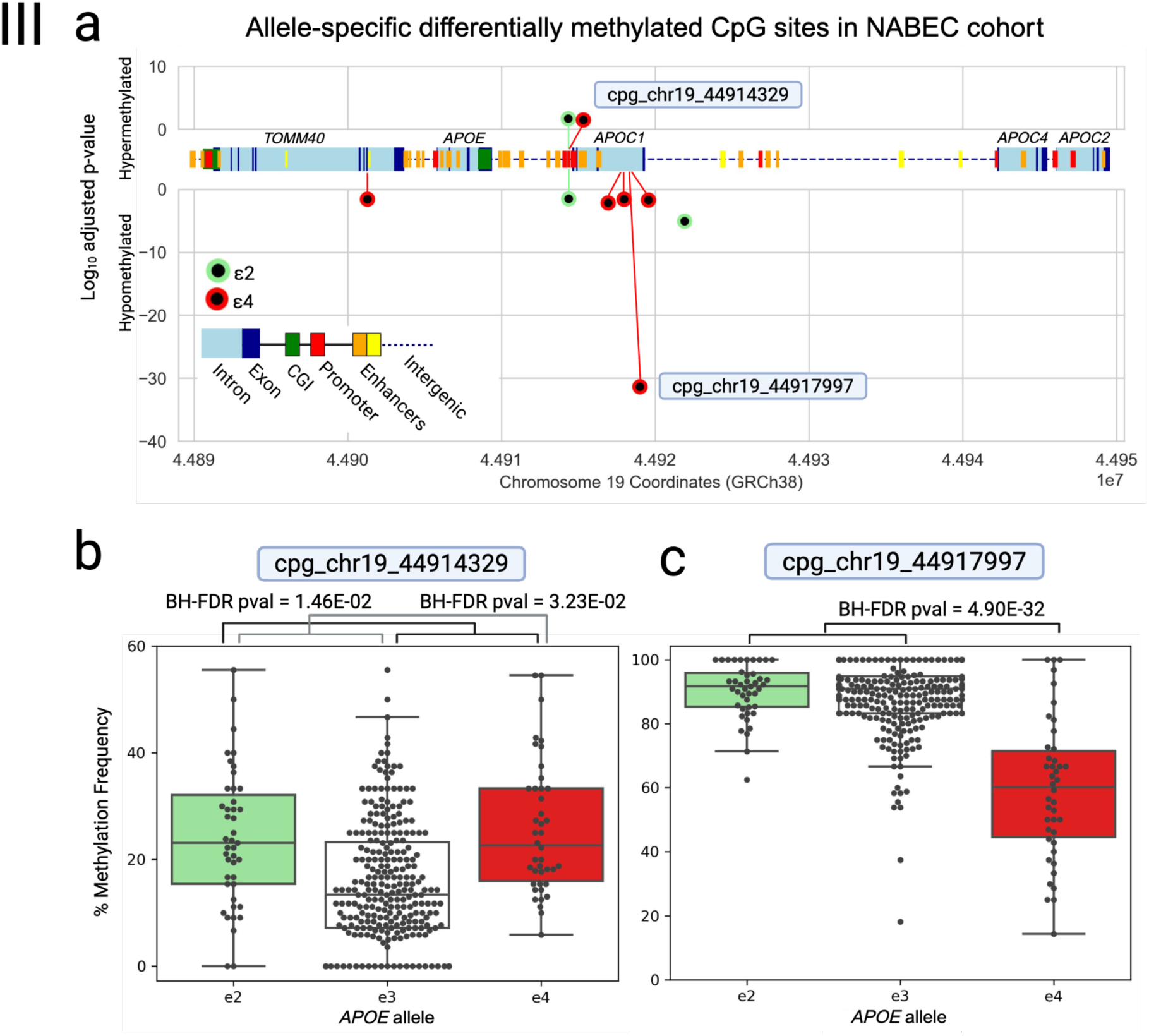
Differentially methylated CpG sites in the *APOE* locus associated with *APOE* ε2 and ε4 alleles in NABEC brain samples. **b)** Association of *APOE*-ε2 (green) and *APOE*-ε4 (red) SNVs with methylation at CpG sites within the *APOE* gene cluster. Each point represents a CpG site that was significantly differentially methylated (BH-FDR-corrected p value < 0.05). The y-axis shows the signed –log_10_(P) value, where positive values indicate hypermethylation and negative values indicate hypomethylation relative to individuals without the respective allele. CpG sites highlighted with labels are shown in the panels below. The middle x-axis provides a schematic of the *APOE* gene cluster region, and the bottom x-axis indicates chromosome 19 genomic coordinates (GRCh38). **b**) Box-and-whisker plot showing methylation levels at CpG site cpg_chr19_44914329, stratified by *APOE* allele status. **c)** Box-and-whisker plot showing methylation levels at CpG site cpg_chr19_44917997, stratified by *APOE* allele status.

**Table 2.**
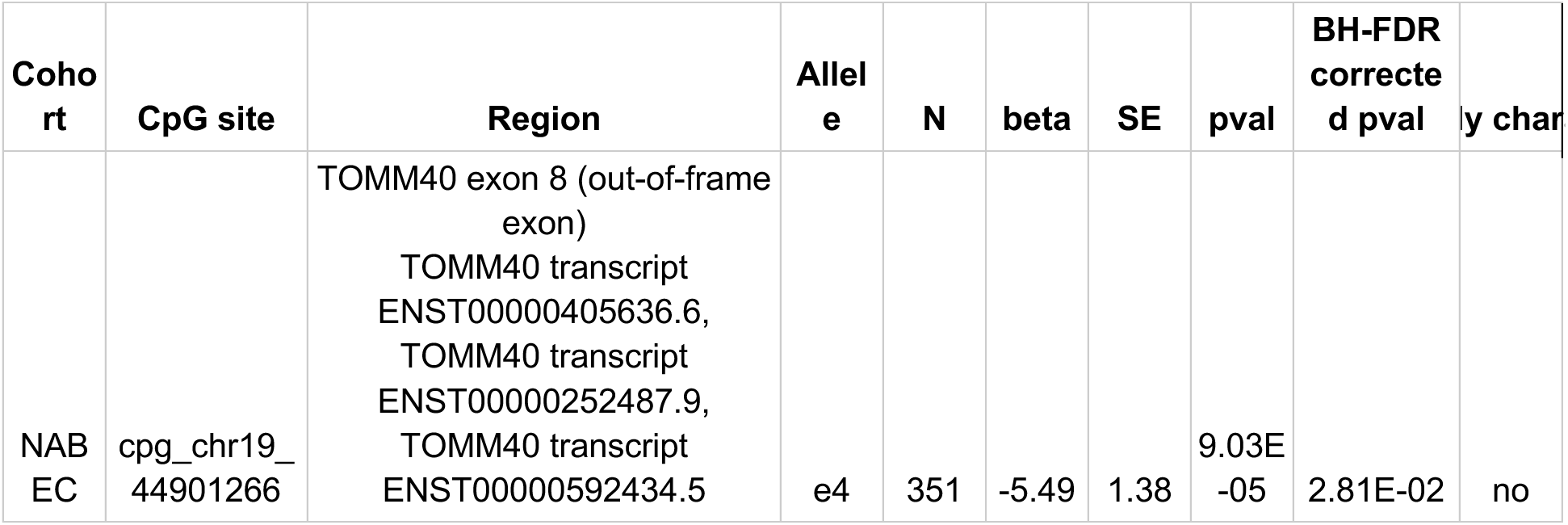
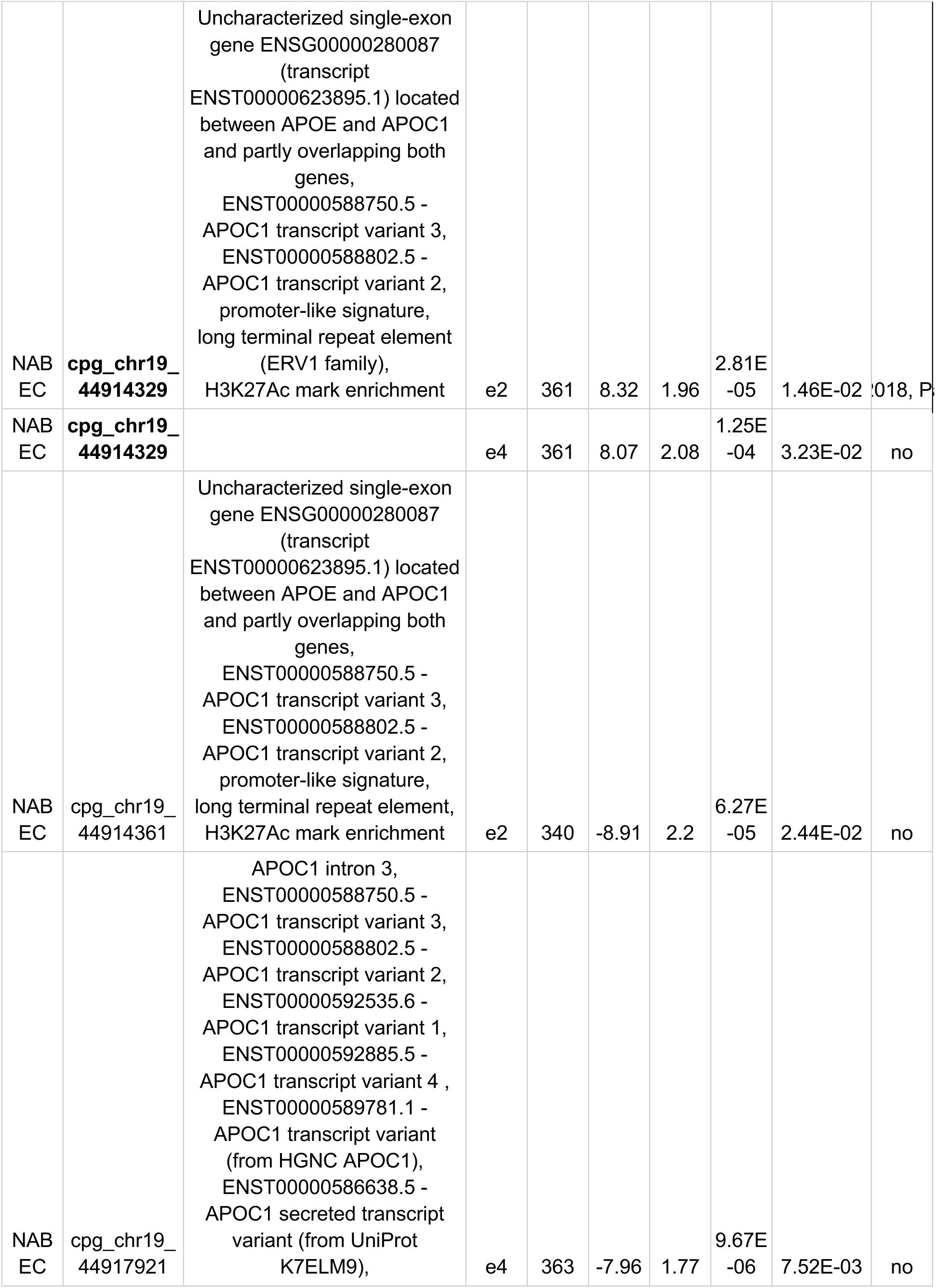
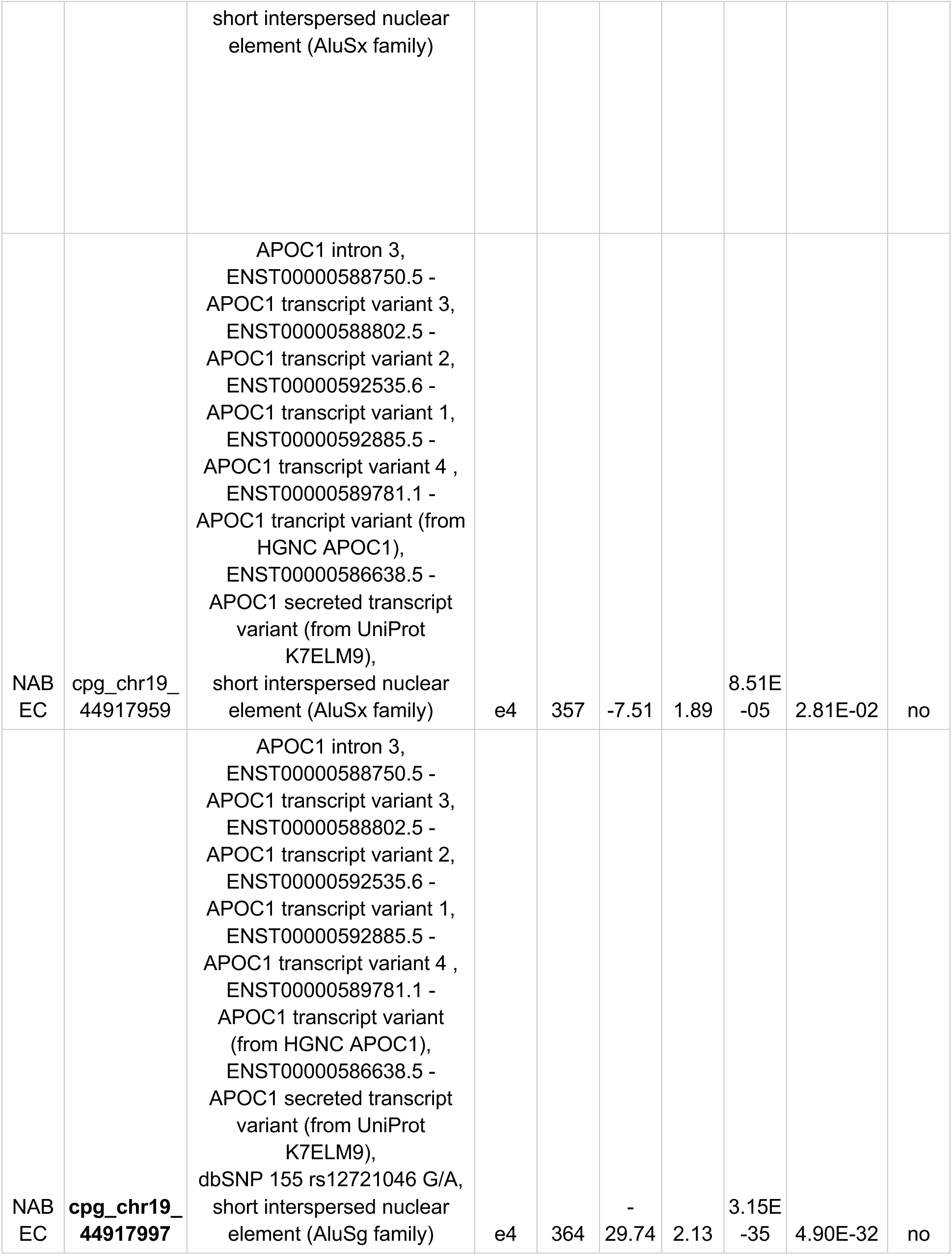
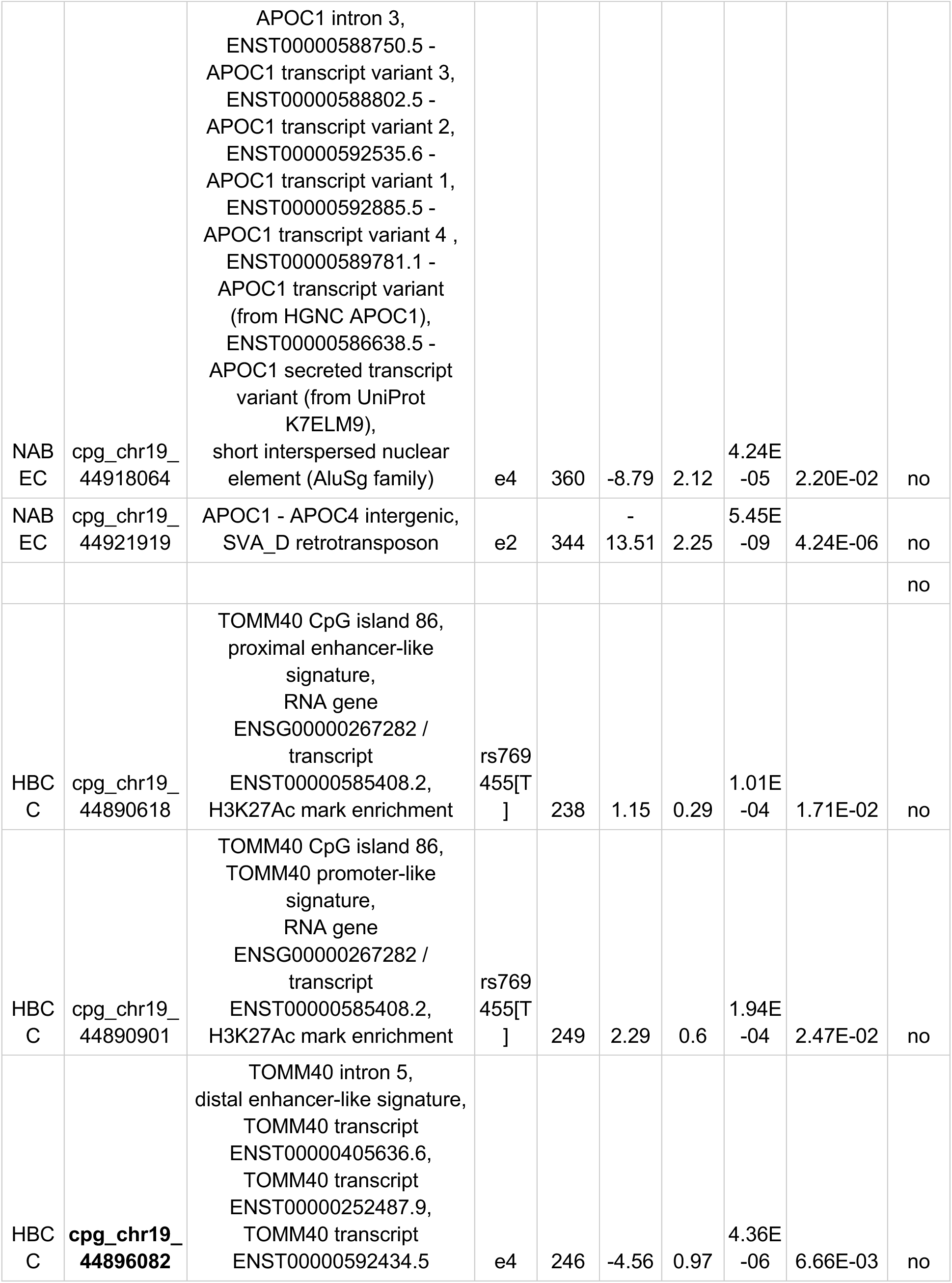
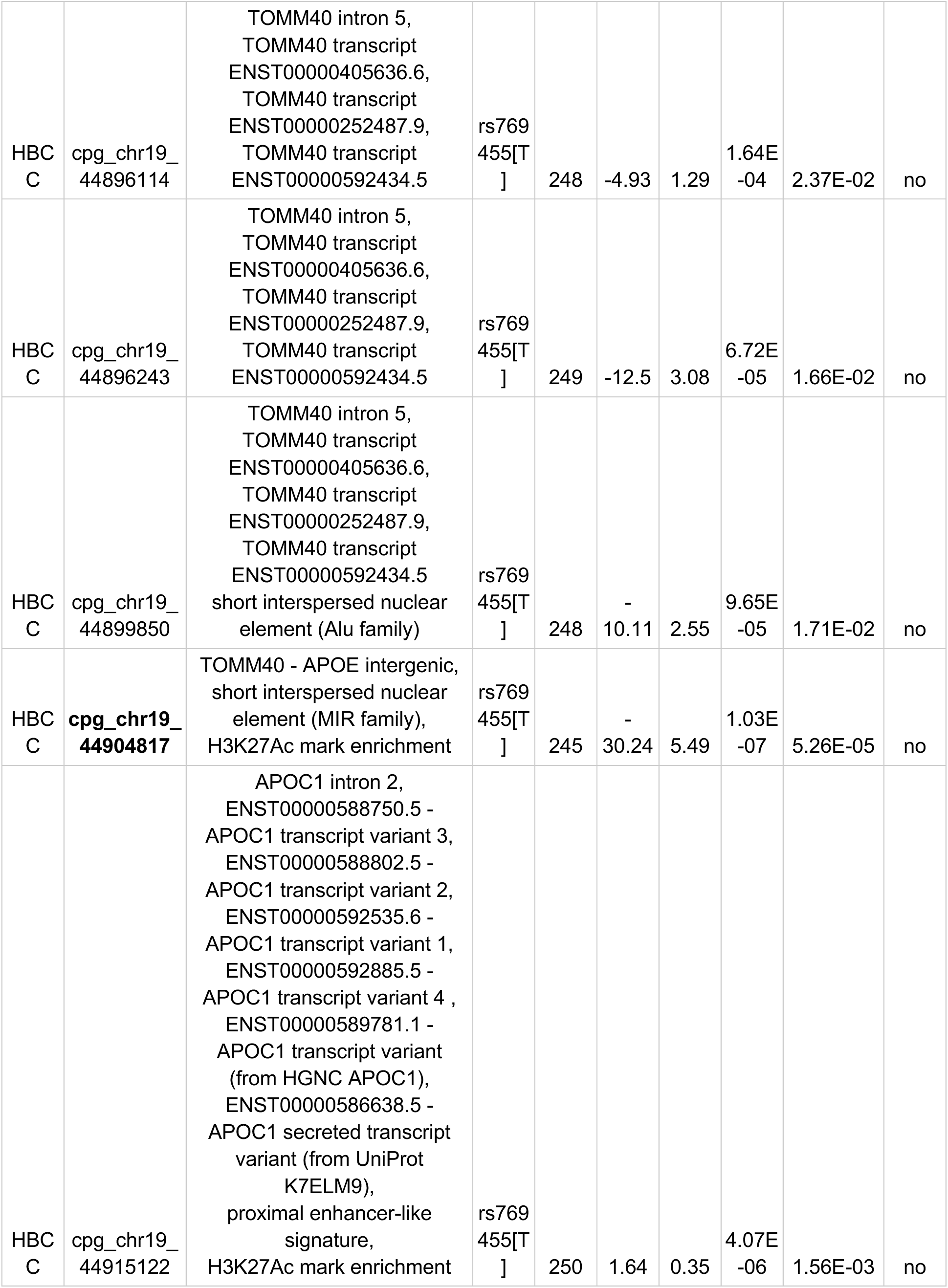
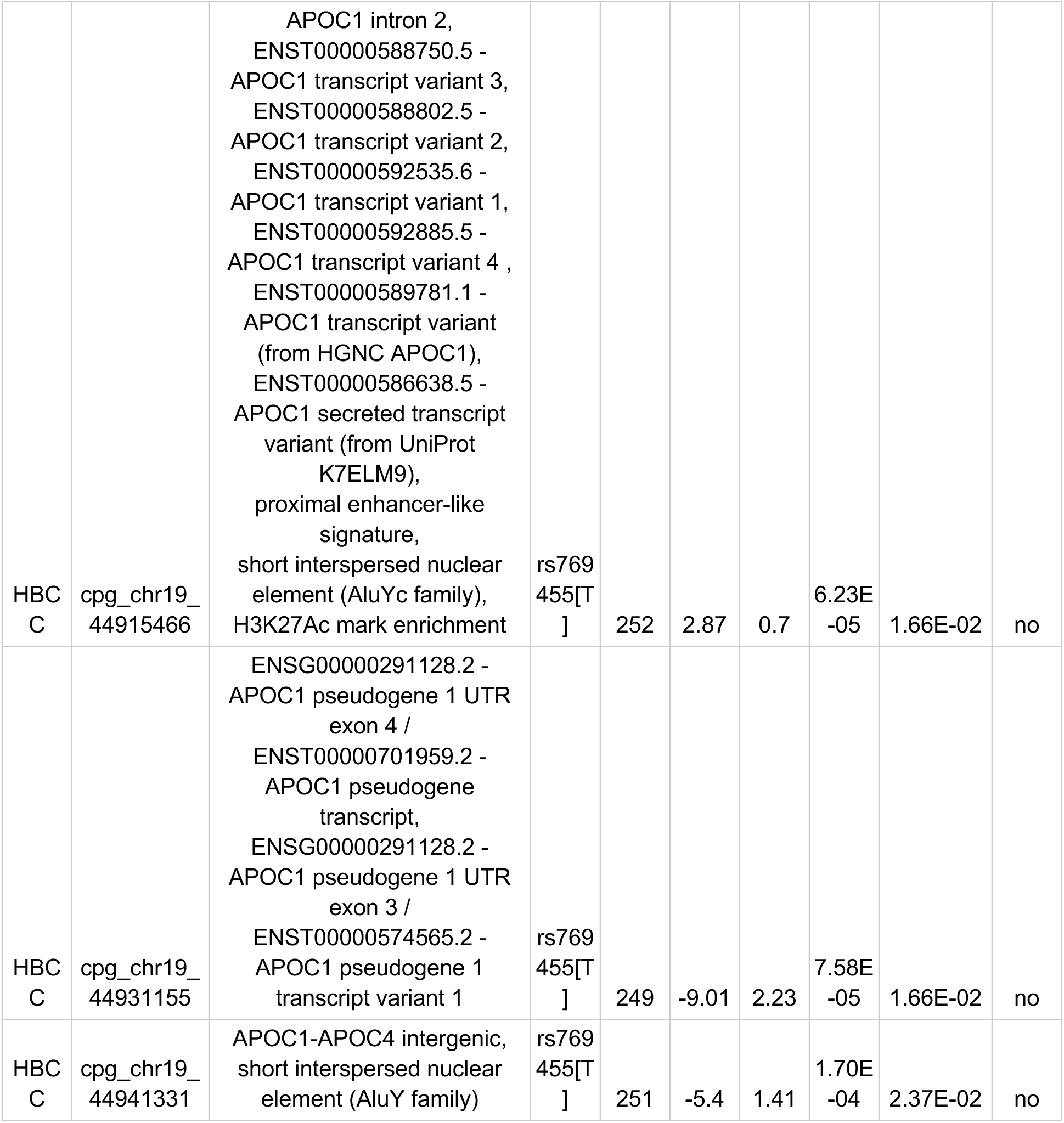
Differentially methylated CpG sites within the *APOE* cluster region in NABEC and HBCC cohorts associated with *APOE* ε2, ε4, or rs769455[T] (BH-FDR-corrected p value < 0.05). SE = Standard Error. Bolded CpG sites are visualized in Figures 3B,C and <u>4B,C</u>.

The most significant methylation difference was observed at cpg_chr19_44917997, where *ε4* alleles were hypomethylated relative to non-*ε4* alleles (BH-FDR-corrected p value = 4.90 × 10^⁻32^, beta =-29.74, SE = 2.13, *P* = 3.15 × 10^-35^; Figure 3C, Table 2). This cpg site is also the site of the *ε4*-linked *APOC1* intronic variant rs12721046[A] which is associated with an increase in AD risk when paired with the *ε*4 allele in non-hispanic white populations ^39–42^. An allele-specific linear regression analysis (described in the methods section) was performed at the rs12721046(G>A) site and yielded a BH-FDR-corrected p value of 1.22 × 10^⁻27^ when *APOE*-*ε*4 was included as a covariate, whereas the *APOE*-*ε*4 allele yielded an insignificant BH-FDR-corrected p value of 0.98 when *APOC1* rs12721046[A] was included as a covariate (Supplemental Figure S4). This indicates that the methylation differences seen at the cpg_chr19_44917997 DMC are more strongly associated with *APOC1* rs12721046[A] allele than the *APOE*-*ε*4 allele, and exemplifies the complexity of epigenetic interactions involving linked variants.

In the HBCC cohort (n = 131, AFR ancestry), 11 CpG sites showed significant allele-specific methylation differences (BH-FDR-corrected p value < 0.05). All 11 CpG sites were novel; none of them were detected as significantly differentially methylated in the NABEC cohort. One site was associated with *ε4* (cpg_chr19_44896082) and the remaining 10 with rs769455[T]. These DMCs were distributed across diverse genomic features, with three in promoter or CpG island regions, four in exons and exon transcripts, six in introns, and four in intergenic regions. Four sites overlapped with short interspersed nuclear elements (SINEs) and four overlapped with predicted enhancer regions (Table 2). The most significant association was observed at cpg_chr19_44904817, where rs769455[A] alleles were hypomethylated compared to non-carrier haplotypes (BH-FDR-corrected *P* = 5.26 × 10^⁻5^, beta =-30.4, SE = 5.49, P = 1.03× 10^-7^; Figure 4B, Table 2).

**Figure 4.**
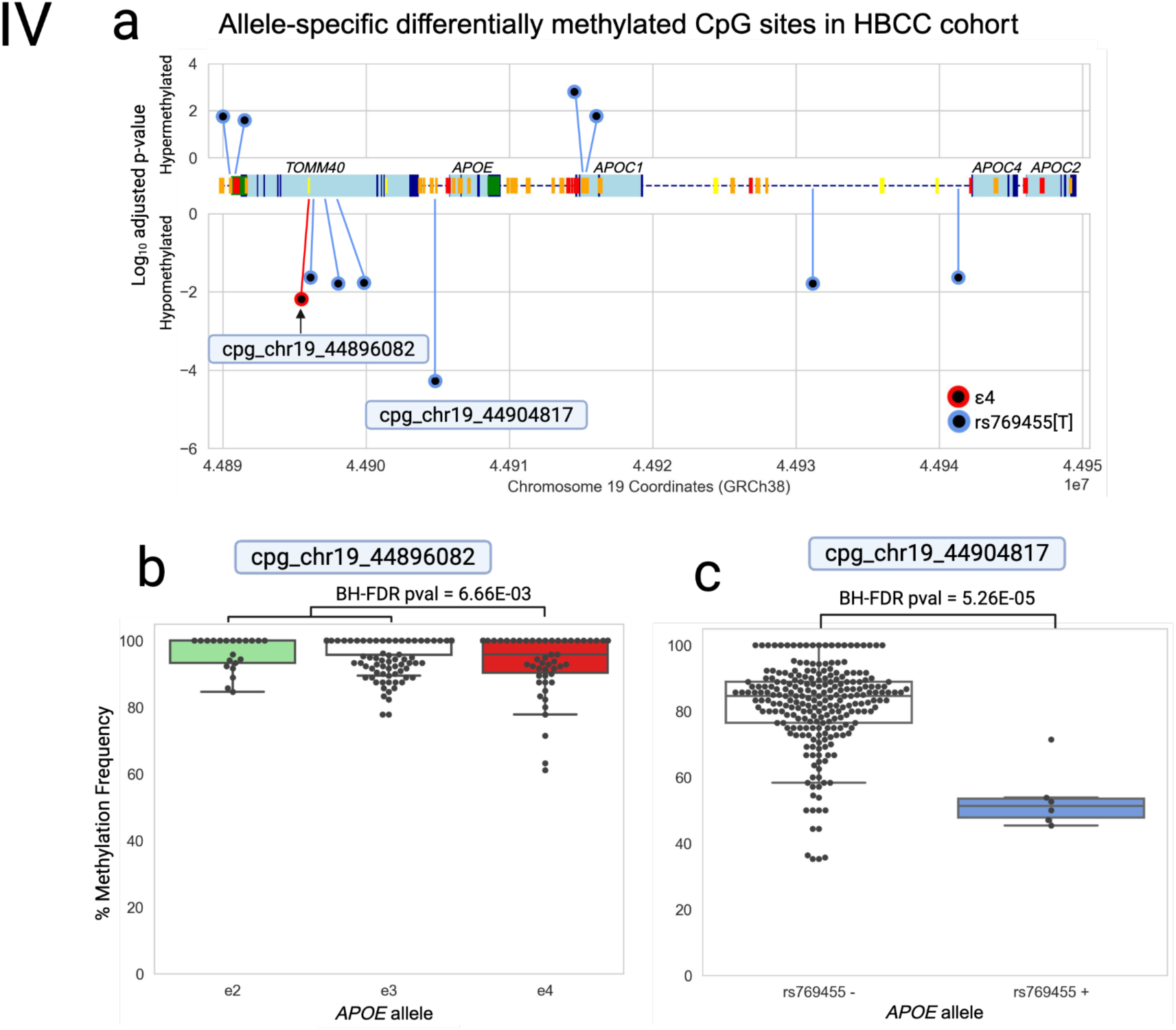
Differentially methylated CpG sites in the APOE locus associated with APOE ε2, ε4, and rs769455[T] alleles in HBCC brain samples. a) Association of *APOE* ε2 (green), ε4 (red) and rs769455[T] alleles with methylation at CpG sites within the *APOE* gene cluster. Each dot represents one of the 11 CpG sites that were deemed significantly differentially methylated (BH-FDR-corrected p value < 0.05) in association with *APOE* ε2 (green dots), ε4 (red dot), or rs769455[T] (blue dots) alleles. The y-axis measures the log_10_ adjusted p-value, with positive numbers indicating hypermethylation of the allele and negative numbers indicating hypomethylation of the allele (in comparison to sample haplotypes without the allele). The labeled CpG sites are graphed in 4b and c. A schematic of the APOE cluster region is shown in the middle x-axis and the bottom x-axis depicts chromosome 19 genome coordinates (reference GRCh38). **b)** Box-and-whisker plot showing methylation frequencies associated with the *APOE* ε4 allele at CpG site cpg_chr19_44896082. **C.** Box-and-whisker plot showing methylation frequencies associated with the *APOE* rs769455 allele in CpG site cpg_chr19_44904817.

A genotype-level methylation analysis was also conducted on the NABEC and HBCC samples in order to highlight the increased sensitivity and resolution of the allele-specific analysis. The genotype-level methylation analysis used the same processing pipeline as the allele-specific methylation analysis except that the input BAM files were not phased. Methylation calls made using unphased data are quantified as the averages of each haplotype; for example, a CpG site that was 0% methylated on haplotype 1 and 100% methylated on haplotype 2 would be interpreted as 50% methylated in the genotype-level analysis. This is illustrated in the representative plots shown in Figure 1B. Since the individual contributions of the different alleles in a heterozygous genotype are unable to be resolved, the effect of an allele on methylation must be quantified by carrier status or dosage effects. For example, instead of examining the direct effect of the ε2 allele type on methylation, one must measure the effect of ε2 dosage (i.e. the effect of no having no ε2 alleles, one ε2 allele, or two ε2 alleles) or ε2 carrier status (i.e. the effect of having an ε2 allele present versus not) on methylation. The confounding effects of heterozygosity can be addressed by including only homozygous samples in the analysis (ex. ε2/ε2, ε3/ε3, and ε4/ε4), however this can greatly reduce sample size and power.

Most methylation analyses rely on bisulfite and array platforms or SRS techniques which lack the ability to resolve haplotypes and assess methylation at the genotype level rather than the allele level. Indeed, all past studies investigating *APOE*-allele-driven DMCs in human brain tissue samples were conducted using unphased data and detected using genotype-level methylation analyses (Supplementary Table S1). To assess the added value of phasing, we compared results from genotype-level and allele-specific methylation analyses in the NABEC cohort. The genotype-level approach identified only three DMCs that were also detected in the allele-specific analysis (cpg_chr19_44914329 (ε2), cpg_chr19_44917997, and cpg_chr19_44921919), two of which were more significant in the allele-specific analysis (Table 2; Supplementary Table S4). None of the allele-specific HBCC methylation differences were found to be significant at the genotype level (Supplementary Table S4). In contrast, the allele-specific linear regression analysis detected 15 differentially methylated CpG sites that were not detected by genotype-level analysis. One example is the NABEC DMC cpg_chr19_44914329, where significant ε4-associated (BH-FDR-corrected p value = 3.23 × 10^⁻2^, beta = 8.07, SE = 2.08, *P* = 1.25 × 10^⁻4^) and ε2-associated (BH-FDR-corrected p value = 1.46 × 10^⁻2^, beta = 8.32, SE = 1.96, *P* = 2.81 × 10^⁻5^) methylation differences were detected (Table 2). This CpG site did not pass significance thresholds in the genotype-level analyses based on homozygous genotype analysis, ε2 carrier status, ε4 dosage or ε4 carrier status (Supplementary Table S4; Supplementary Figure 5B-F), and serves as just one example of the many modestly significant methylation differences that were missed by the genotype-level methylation analysis.

Interestingly, cpg_chr19_44914329 was also the only DMC that was identified by previous studies looking at *APOE* allele related and/or AD-related methylation differences in human brain or blood samples. A study by Shao et al. (2018) used an Illumina Infinium HumanMethylation450 BeadChip array to look for AD-related methylation differences in 54 CpG sites in human brain and blood samples (Supplementary Table S1). A multivariate analysis revealed that the *APOC1* promoter CpG site cg2327011 (which is also cpg_chr19_44914329) was differentially methylated in control and AD blood samples in the study’s replication cohort data from Gene Expression Omnibus (*P* = 0.019), however the significance disappeared after Holm correction for 54 CpG sites (BH-FDR corrected p value = 1.002).

A more recent study by Panitch et al. (2024) used an Illumina Infinium HumanMethylation450 BeadChip assay to measure genome-wide methylation differences associated with AD and/or *APOE* genotype in human blood and brain tissue samples (Supplementary Table S1). They found that cg23270113 (which is also cpg_chr19_44914329) was significantly hypomethylated in *APOE* ε4 carriers compared to non-carriers in control blood samples (T=-6.40, P=1.5 x 10^-10^)^24^. Hypomethylation at this site was also associated with poor memory performance in *APOE* ε4 non-carriers, while hypermethylation was associated with worse performance on cognitive performance tests in ε4 carriers (Supplementary Table S1).

To explore whether *APOE* allele-driven methylation differences affect gene expression, we performed a linear regression analysis using bulk RNA-seq data that was available for all of the NABEC cohort samples and 86 HBCC cohorts ^36^. The expression of each detectable gene in the *APOE* cluster region (*TOMM40, APOE, APOC1, APOC4-APOC2*) was tested for associations with each *APOE* allele (ε2, ε4 and rs769455) with no significant associations observed (Supplementary Table S5).

## Discussion

This study represents the most comprehensive analysis to date of allele specific DNA methylation at the *APOE* locus in the human brain, enabled by high-accuracy long-read sequencing. By leveraging publicly available data from the NIH CARD Long-Read Sequencing Initiative, we analyzed 332 postmortem brain samples from individuals of EUR and AFR ancestry, generating phased methylomes that captured a level of epigenetic resolution previously inaccessible with array and bisulfite based techniques. Our findings highlight the power of ONT LRS for identifying allele-specific methylation patterns at functionally important loci. Compared to Illumina-based methylation arrays, long-read sequencing enabled ∼33-fold greater CpG site detection across the *APOE* cluster region, extending coverage beyond promoter and exonic regions into intronic, intergenic, and regulatory elements. This expanded view of the brain methylome facilitated the discovery of 18 novel DMCs associated with *APOE* ε2, ε4, and rs769455 alleles. These DMCs were enriched in regions with known regulatory potential, including SINE elements, promoters, enhancers, and CpG islands, providing a more complete map of epigenetic variation that may contribute to AD risk.

One of the few CpG sites within the *APOE* locus that overlapped with an Illumina Infinium HumanMethylation450 BeadChip array probe was cpg_chr19_44914329, which also represents the only CpG detected in our study that has been reported in previous methylation analyses. Notably, Panitch et al. (2024) found that this site was hypomethylated in peripheral blood samples from *APOE* ε4 carriers compared to non-carriers. In contrast, our brain-based analysis revealed that cpg_chr19_44914329 was hypermethylated on ε4 alleles relative to ε3 and ε2 alleles. This discrepancy could reflect tissue-specific methylation patterns, which are well-documented in the literature ^43–45^ and emphasize the importance of studying methylation within disease-relevant tissues such as the brain.

Previous studies investigating allele-specific methylation at the *APOE* locus have primarily focused on the ε4 allele, given its strong association with AD. In contrast, the ε2 allele remains understudied, largely due to its lower frequency in the population, which limits statistical power in many cohorts. Leveraging phased data and an allele-specific analytical framework, we were able to isolate 73 ε2 alleles (48 from NABEC and 25 from HBCC) and identify three ε2-associated DMCs, two of which were located in putative regulatory elements. These findings provide some of the first evidence of ε2-specific methylation patterns in the human brain and offer new insights into the epigenetic mechanisms that may underlie the allele’s protective role in AD.

Despite these advances, several limitations should be considered. First, NABEC and HBCC samples were sequenced using different ONT chemistries (R9 and R10, respectively), which may introduce technical variation in variant and methylation calling. Subtle methylation differences detected by an allele-specific analyses are more likely to be affected by chemistry-related methylation calling differences, which make it difficult to combine cohorts in analysis ^35^. Second, sample size remains a limiting factor, particularly in the African ancestry cohort, where the number of individuals carrying ε2 or rs769455 alleles was relatively small. This limited our power to detect allele-specific methylation differences and prevented genotype-based modeling for certain variants. Expanding representation of ancestrally different populations will be critical to fully capture the regulatory landscape of *APOE* and its functional variants. Third, we evaluated potential gene expression effects of differentially methylated CpG sites using bulk RNA-seq data from the NABEC and HBCC cohorts. However, these data were generated using SRS, which lacks the resolution to capture full-length transcript isoforms or alternative splicing events. Additionally, short-read RNA-seq has limited ability to resolve allele-specific expression, particularly in complex, polymorphic regions such as the *APOE* locus. Although no significant eQTLs were identified in this study, this may reflect the limited resolution of SRS data rather than a true absence of regulatory effects. Future studies using long-read RNA sequencing will be critical for resolving isoform diversity and quantifying allele-specific transcript expression, enabling a more accurate assessment of the functional consequences of methylation variation. Finally, all samples in this study were derived from neurologically healthy control brains. The lack of observed expression changes may therefore reflect the absence of disease-related transcriptional dysregulation. Investigating these methylation patterns, and their regulatory impacts, in Alzheimer’s disease and related dementias (ADRD) cohorts will be essential for understanding their potential contribution to disease risk and progression.

Looking ahead, this work provides a strong foundation for future functional studies aimed at elucidating how *APOE* allele-specific methylation modulates gene regulation, lipid metabolism, and AD pathogenesis. As part of the NIH’s ongoing CARD Long-Read Sequencing Initiative, thousands of additional brain samples, including individuals affected by ADRDs, are being sequenced using improved R10 chemistry with higher throughput and accuracy. Expanding this approach to include disease cohorts and integrating with multi-omic datasets, such as long-read transcriptomics, chromatin accessibility, and single-cell data, will be key to fully resolving the regulatory mechanisms through which *APOE* variation influences disease risk across diverse populations. In summary, our study highlights the unique advantages of LRS for resolving the complexity of *APOE* haplotypes and their epigenetic architecture in the human brain. These findings offer new insight into the molecular mechanisms linking *APOE* variation to AD risk and pave the way for more precise, ancestry-aware models of disease biology.

## Methods

### Cohort information

We utilized existing ONT whole genome sequencing (WGS) data from the North American Brain Expression Consortium (NABEC) and the Human Brain Collection Core (HBCC) cohorts ^36^. The NABEC cohort consists of 206 prefrontal brain tissue samples from individuals of European ancestry and the HBCC cohort consists of 142 prefrontal cortex brain tissue samples from individuals of predominantly African and African-admixed ancestries. Ancestries were determined genetically using the GenoTools package ^46^. All samples were derived from individuals with no known neurological diseases or clinical history of cognitive impairment. Four of the NABEC samples and ten of the HBCC samples were excluded from the study due to receiving non-European (NABEC) or non-African or African admixed (HBCC) genetic ancestry estimations by the GenoTools package ^46^, and one sample from each cohort was excluded due to having low read coverage across the *APOE* gene cluster region. Additional information about cohort demographics and sequencing statistics can be found in Supplementary Tables S2 and S3.

### Data source and processing

We accessed preprocessed, aligned BAM files previously generated and publicly released by Billingsley et al. (2024). Detailed protocols for DNA extraction, library preparation, and sequencing are available on the protocols.io platform ^47,48^. Briefly, DNA was extracted from approximately 40 mg of homogenized frozen frontal cortex tissue and sheared to a target fragment size of 30 kb. NABEC samples were prepared using the ONT SQK-LSK110 kit and sequenced on R9.4.1 flow cells, while HBCC samples were prepared using the SQK-LSK114 kit and sequenced on R10.4.1 flow cells. All sequencing was performed on PromethION flow cells (FLO-PRO002), with 2–3 rounds of loading to achieve minimum yield thresholds. The NABEC samples were sequenced using MinKNOW versions 22.03.2 through 22.10.7 and basecalled with Guppy v6.1.2, while the HBCC samples were sequenced with MinKNOW v22.10.7 and basecalled using Guppy v6.3.8. Read alignment to the GRCh38 reference genome was performed using minimap2 (v2.23-r1111))^49^.

### Haplotype phasing and methylation calling

Reads were aligned to the GRCh38 reference genome using minimap2 followed by haplotype phasing and downstream methylation analysis. Reads from chromosome 19 were subset from each sample, and phasing was conducted using PEPPER-Margin-DeepVariant (ONT_R9 model) for NABEC samples and DeepVariant (ONT_R104 model) for HBCC samples ^50,51,52^. The--dv_sort_by_haplotypes parameter was applied to produce haplotype-resolved BAMs. DeepVariant output gVCFs were used to determine each sample’s *APOE* genotype. Original unphased BAMs were retained to compare phased versus unphased methylation profiles.

Methylation calling was carried out on both phased and unphased BAM files using modkit pileup (https://github.com/nanoporetech/modkit), with the--combine-strands and--ignore h options enabled to extract CpG-level methylation frequencies. This pipeline enabled us to quantify allele-specific methylation differences across the *APOE* locus. A schematic overview of the haplotype-specific methylation analysis workflow is presented in Figure 1A.

### Selection and classification of *APOE*alleles

We selected single nucleotide variants for this study that were: (i) located within the *APOE* gene cluster; (ii) had a minor allele frequency (MAF) > 0.01 in individuals of European (EUR) and/or African and African admixed (AFR) ancestry; (iii) were classified as protective or pathogenic (CADD PHRED > 20); and (iv) had previously reported being significantly associated with AD risk. The following variants met these criteria: *APOE* ε2 (rs7412(C>T)), ε4 (rs429358(T>C)), and rs769455(C>T).

### Dendrogram generation

Dendrograms were generated using the pipeline described by Chang et al (2021) (https://bioinformaticsworkbook.org/phylogenetics/FastTree.html#gsc.tab=0) with several adaptations. Briefly, *bedtools* ^53^ was used to extract reads in the *APOE* gene region (chr19:44889556–44953378; hg38) from both phased and unphased BAM files in the NABEC and HBCC cohorts. *Samtools* ^54^ was then used to split the phased reads by haplotype using the HP tag and each haplotype was treated as an individual sample (i.e. HBCC_81925_FTX becomes HBCC_81925_FTX_H1 and HBCC_81925_FTX_H2). *Bedtools genomecov* was used to filter and include haplotypes with ≥10× coverage for the phased dataset and ≥20× coverage for the unphased dataset.

Each BAM file was then converted into its consensus FASTA sequences using *samtools consensus*. A NABEC sample that had been sequenced with both R9 and R10 chemistries was included to assess whether dendrogram separations were driven by chemistry differences. A FASTA file of the ancestral chimpanzee *APOE* sequence was also included to root the tree^38^. All FASTA sequences were concatenated and aligned using the *Mafft* ^55^ package with default parameters, and phylogenetic trees were constructed using the *FastTree* package ^56^. The resulting Newick files were visualized using *FigTree* (https://github.com/rambaut/figtree/) with aligned tip labels, ordered nodes, and coloring based on allele-type.

### Methylation linear regression analysis

Allele-specific differentially methylated CpG sites in the *APOE* cluster region were identified using an ordinary least squares (OLS) linear regression model with the presence or absence of the SNV as the independent variable and the modkit-determined phased CpG methylation frequencies as the dependent variable. Covariates included age, sex, post-mortem interval (PMI), brain bank sources (‘SH’, “KEN‘’, and ‘UMARY’ for NABEC), and genetic and methylation principal components (PCs). Genetic PCs were generated using PLINK (v1.9), with the exclusion of variants with a minor allele frequency <0.05, genotyping rate <0.95, and Hardy-Weinberg equilibrium p value > 0.0001. Methylation PCs were calculated using genome-wide, haplotype-specific CpG site methylation frequency values generated by modkit. The number of PCs included in the analysis was determined using scree plots of the cumulative variance, with the included PCs capturing below 70% of the variance for genetic PCs or, or the first five PCs if cumulative variance is below 70% for the first 20 PCs. This criteria resulted in the inclusion of the first 14 genetic PCs and first 5 methylation PCs for both NABEC and HBCC cohorts. Methylation PCs for haplotype 1 and 2 were highly correlated, so only haplotype 1 PCs were used in the analysis. A separate linear regression analysis was run using modkit-determined unphased CpG methylation frequencies as the dependent variable in order to see which CpG sites would be detected with unphased input. Genetic PCs were reused from the allele-specific analysis and methylation PCs were recalculated using genome-wide unphased CpG site methylation frequency values. The first 14 genetic PCs and first 5 methylation PCs were included for both NABEC and HBCC cohorts in the unphased analysis.

More details about the files and file formats are located on the NIH-CARD Github page (https://github.com/NIH-CARD/CARDlongread_data_standardization) and the linear regression scripts are located at https://github.com/rgenner/Allele_specific_meth_scripts. CpG sites with a FDR-adjusted/Benjamini-Hochberg (BH) corrected p value < 0.05 were considered significant.

### Expression linear regression analysis

Bulk RNA-seq data for all of the NABEC samples^57^ and 86 of the HBCC samples (https://nda.nih.gov/edit_collection.html?id=3151) previously processed by Billingsley et al (2024) ^36^ using Salmon ^58^ with the filtered Gencode v43 human transcriptome index. In the NABEC samples, 26,778 genes were considered to be well-detected (missingness under 0.25) including the *TOMM40*, *APOE* and *APOC1* genes. In the 86 HBCC samples, 26,679 genes were considered to be well-detected, including the *TOMM40, APOE, APOC1 and APOC4-APOC2* genes. Expression data was quantile transformed and scaled from 0 to 1. *APOE* SNV-related expression differences in the detected *APOE* cluster genes were tested for using an OLS regression model with the presence or absence of the SNV as the independent variable and the standardized gene expression data as the dependent variable. Covariates included age, sex, post-mortem interval (PMI), brain bank sources (‘SH’, “KEN‘’, and ‘UMARY’ for NABEC), and genetic and expression principal components. Genetic PCs were generated using PLINK (v1.9), with the exclusion of variants with a minor allele frequency <0.05, genotyping rate <0.95, and Hardy-Weinberg equilibrium p value > 0.0001. Gene expression PCs were calculated using the sklearn.decomposition PCA module. The number of PCs to include in the analysis was determined using scree plots (the first 14 genetic PCs and first five expression PCs were included for both NABEC and HBCC cohorts). Gene expression differences with a FDR-adjusted/Benjamini-Hochberg (BH) corrected p value < 0.05 were considered significant.

### Reference methylation data from the ROSMAP Study

DNA methylation data shown in Figure 2a and Supplementary Figure 1 were derived from the Religious Orders Study and the Rush Memory and Aging Project (ROSMAP) cohorts ^37,59^. We analyzed data from the previously produced Illumina HumanMethylation450 BeadChip array, generated from dorsolateral prefrontal cortex tissue of 697 participants, including 417 individuals with AD and 280 cognitively normal controls.

## Acknowledgments

This work was supported in part by the Intramural Research Program of the National Institutes of Health including: the Center for Alzheimer’s and Related Dementias, within the Intramural Research Program of the National Institute on Aging and the National Institute of Neurological Disorders and Stroke. This work utilized the computational resources of the NIH HPC Biowulf cluster (https://hpc.nih.gov).

## Ethics declaration / Competing Interests

The NABEC and HBCC sequencing data used in these analyses was derived from brain tissue samples that were processed and sequenced for a previous study by Billingsley et al. (2024) as part of the CARD Long-read Initiative. This study obtained all brain tissue samples using proper approvals, protocols and documentation. The NIH considered research using post-mortem material as nonhuman subject research and therefore no additional institutional review board approval was required.

This research was supported in part by the Intramural Research Program of the NIH, National Institute on Aging (NIA), National Institutes of Health, Department of Health and Human Services; project number ZO1 AG000534. This article is a US Government work. It is not subject to copyright under 17 USC bioRxiv preprint doi: https://doi.org/10.1101/2024.12.16.628723; this version posted December 18, 2024. This work utilized the computational resources of the NIH HPC Biowulf cluster (https://hpc.nih.gov). Some authors’ participation in this project was part of a competitive contract awarded to DataTecnica LLC by the National Institutes of Health to support open science research. M.A.N. also owns stock in Character Bio Inc. and Neuron23 Inc.

## Data availability

Human brain sequencing datasets are under controlled access and require a dbGap application (phs001300.v4) (phs000979.v4) and are then available through the restricted AnVIL workspace (add link when live).

## Code availability

The code used to process and analyze the data for this study is publicly available at https://github.com/rgenner/Allele_specific_meth_scripts. This repository includes scripts designed to conduct an allele-specific, CpG-site specific linear regression analysis given a modkit bed file for a specific region and sample haplotype designations. The scripts used to generate all main and supplemental figures are also included. Additional details about the computational tools and parameters used in this study are described in the Methods section of the manuscript.

## Supplementary Figures

**Supplementary Figure 1.**
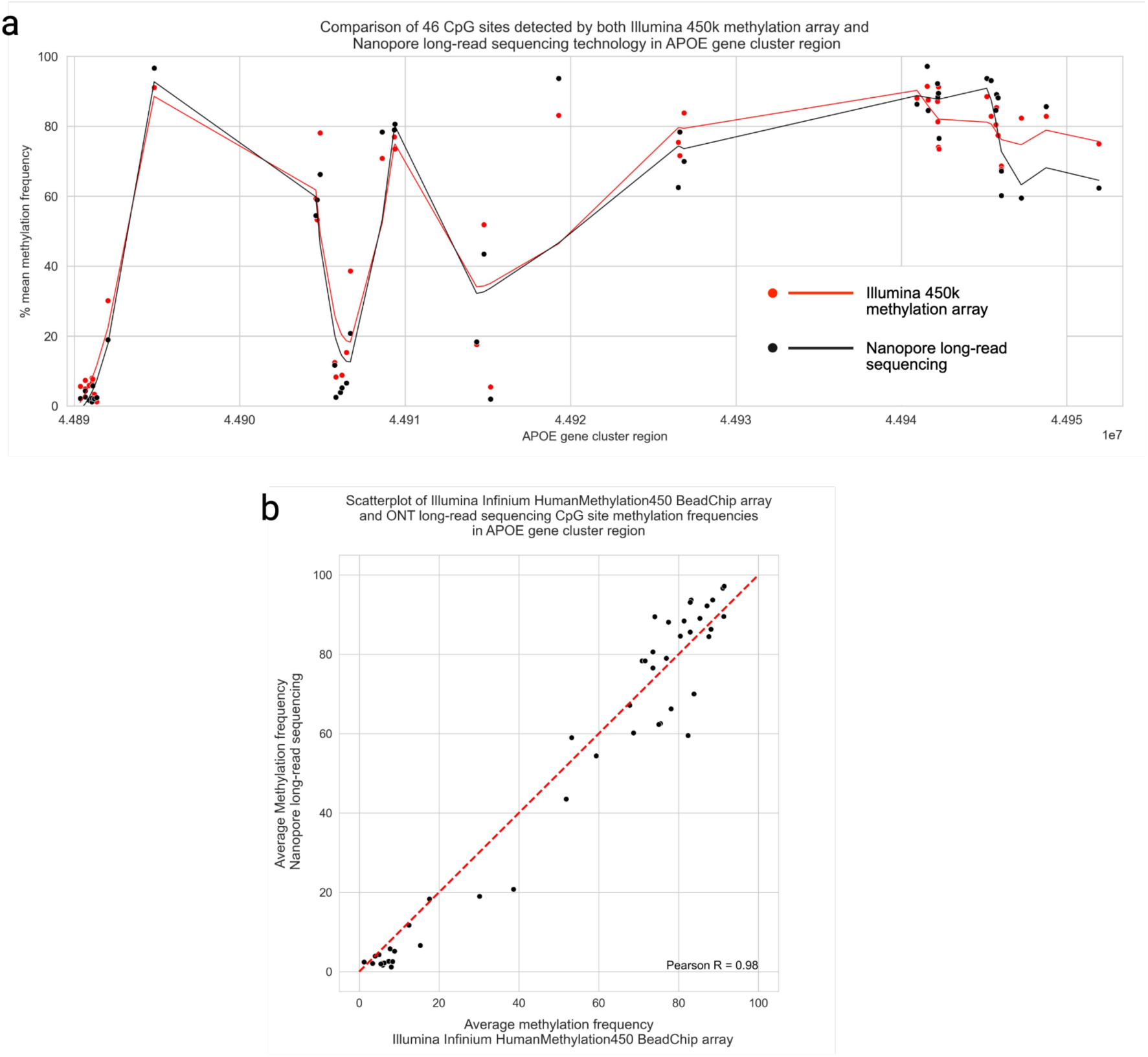
Comparison of the 46 CpG sites detected by both platforms. **a)** Mean methylation frequencies of the 46 CpG sites detected in two separate EUR-ancestry brain tissue cohorts sequenced by the Illumina Infinium HumanMethylation450 BeadChip array (red) and the ONT long-read sequencing technology (black) within the *APOE* cluster region (*TOMM40, APOE, APOC1,* and *APOC4-APOC2* genes; hg38 coordinates chr19:44889556-44953378). **b)** Scatter plot of the Illumina Infinium HumanMethylation450 BeadChip array and the long-read sequenced CpG site methylation frequencies in the *APOE* cluster region. Pearson R = 0.98, P-value = 3.506e-33.

**Supplementary Figure 2.**
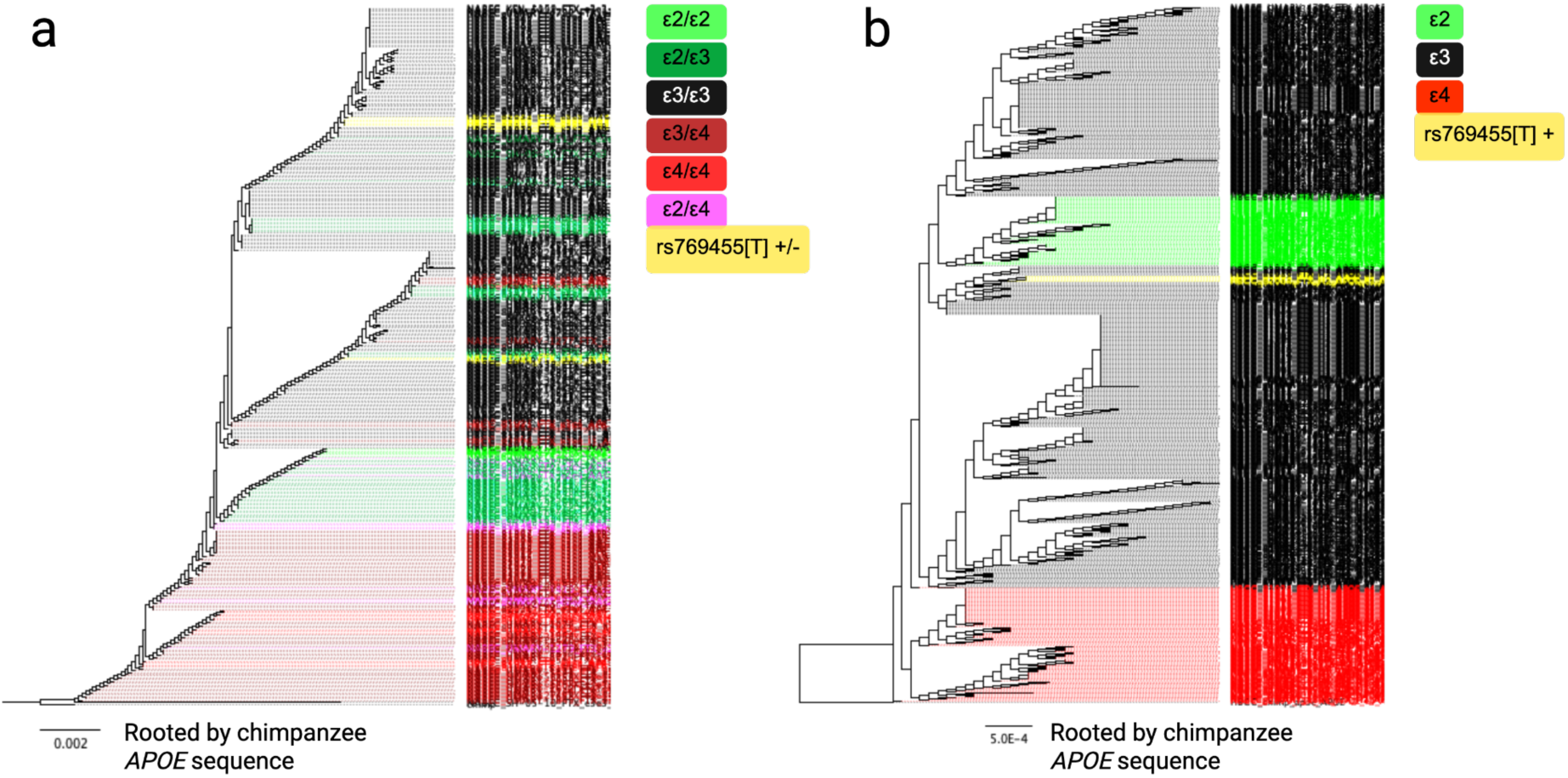
Dendrograms depictions of *APOE* allele separation at the allele and genotype levels. **a)** Dendrograms of the *APOE* gene region (chr19:44905791-44909393, hg38) generated from 332 unphased genotypes from the NABEC and HBCC cohorts. Branches are colored by *APOE* genotype. Only basecalls with >20× coverage were included. **b)** Dendrogram of the *APOE* region generated from 664 phased haplotypes from the NABEC and HBCC cohorts. Branches are colored by allele type: *APOE*-ε2 in green, *APOE*-ε4 in red, and rs769455[T] in yellow. Only positions with > 10× coverage were included. In both panels, each branch represents the aligned consensus FASTA sequence for an individual *APOE* genotype (a) or haplotype (b). Dendrograms were rooted using the ancestral chimpanzee *APOE* sequence. See Methods for full details on dendrogram generation.

**Supplementary Figure 3.**
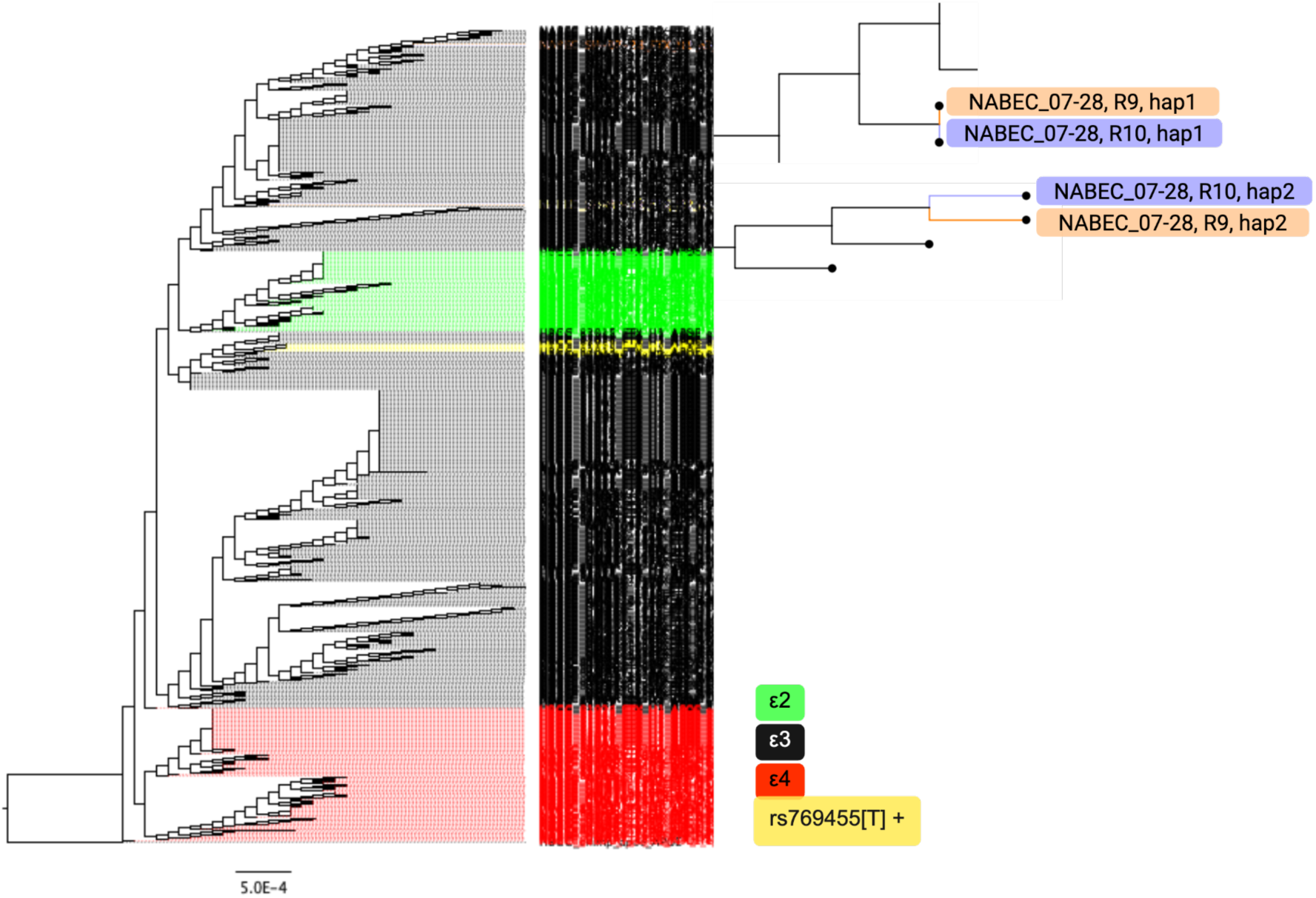
Benchmarking the effect of Nanopore flow cell chemistry on phased allele separation in a dendrogram. A copy of the dendrogram featured Supplementary Figure 2 that has been expanded to show one NABEC sample (NABEC_07-28) that was sequenced with both R9 (orange) and R10 (purple) chemistries. The branches containing each sample haplotype have been magnified on the right side of the dendrogram.

**Supplementary Figure 4.**
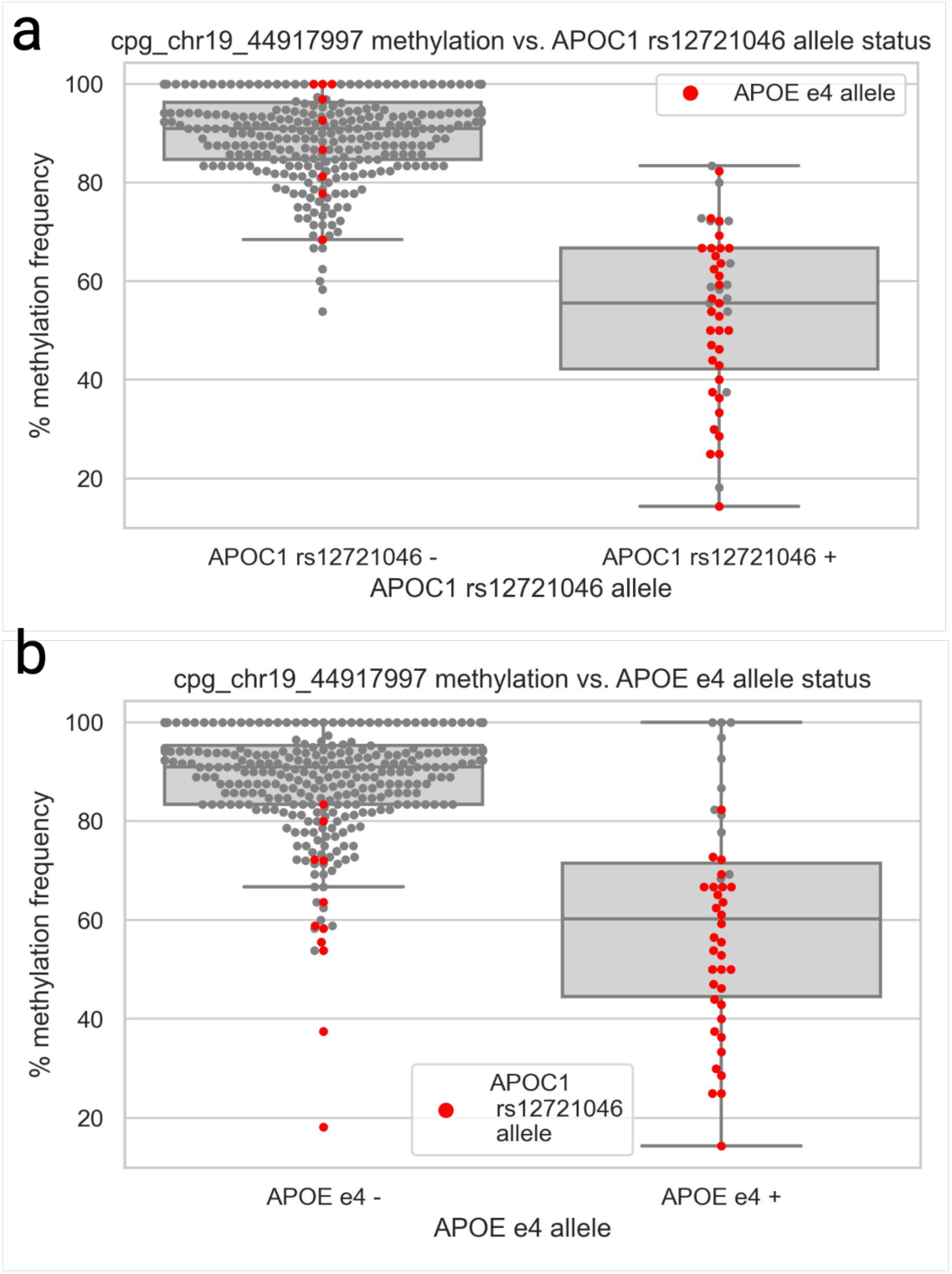
Assessing *APOE* e4 and *APOC1* rs12721046[A] allele effects on methylation frequency differences at NABEC CpG site cpg_chr19_44917997. a) A box-and-whisker plot showing the methylation frequency difference at NABEC CpG site cpg_chr19_44917997 when stratified by the *APOC1* rs12721046[A] allele (with the *APOE*-ε4 alleles highlighted in red and included as a covariate in the linear regression analysis). **b)** A box-and-whisker plot showing the methylation frequency difference at NABEC CpG site cpg_chr19_44917997 when stratified by the *APOE*-ε4 allele (with the *APOC1* rs12721046[A] + alleles highlighted in red and included as covariate in linear regression analysis). The dots on the left half of the box plots depict sample haplotypes that do not have the allele of interest (denoted by “-”) and the right half show samples that do have the allele of interest (denoted by “+”).

**Supplementary Figure 5.**
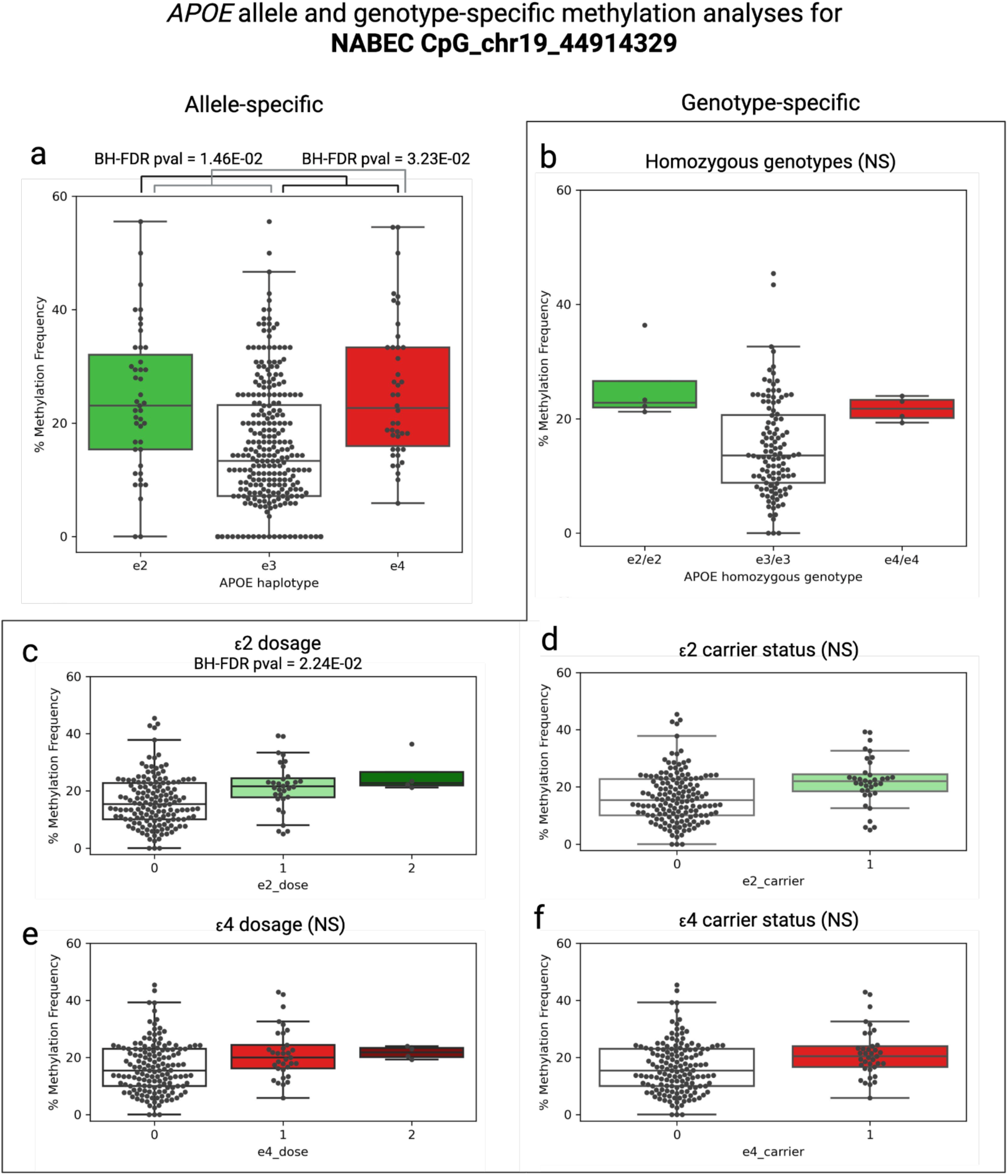
Box-and-whisker plots of the ε4 *APOE* allele-specific methylation frequencies for NABEC CpG site cpg_chr19_44914329 as determined by allele-specific and genotype-level methylation analyses. **a)** The boxplot of allele-specific methylation analysis results shown in Figure 3b. **b-f)** Genotype-level methylation analyses based on homozygous genotypes (b), ε2 dosage and carrier status (c, d) and ε4 dosage and carrier status (e, f). NS = BH-FDR corrected p value was not significant.

## Supplementary Tables

**Table S1.** Summary table of previous studies of *APOE* methylation in brain and blood tissue samples including the number and type of samples analyzed, methylation sequencing technique used, number of CpG sites identified and where they were located in the genome.

**Table S2:** Summary of cohort demographics including the total number samples, number of male and female samples, mean age, age range, ancestry, diagnosis, sample type, sequencing coverage and sequencing N50.

**Table S3:** Haplotype-specific sequencing statistics for each NABEC and HBCC sample used in the study including the mean and median sequencing lengths, N50, number of reads, and mean coverage.

**Table S4:** Significant allele-specific differentially methylated CpG sites analyzed using genotype-level methylation analysis methods.

**Table S5:** Linear regression results of *APOE* cluster region gene expression associated with *APOE* alleles.

## References

1. Strittmatter, W. J. & Roses, A. D. Apolipoprotein E and Alzheimer disease. Proc. Natl. Acad. Sci. U. S. A. 92, 4725–4727 (1995).

2. Mahley, R. W., Weisgraber, K. H. & Huang, Y. Apolipoprotein E: structure determines function, from atherosclerosis to Alzheimer’s disease to AIDS. J. Lipid Res. 50 Suppl, S183–8 (2009).

3. Apolipoprotein, E. Alzheimer disease: risk, mechanisms and therapy. Nature Reviews Neurology.

4. Mahley, R. W. Apolipoprotein E: cholesterol transport protein with expanding role in cell biology. Science 240, 622–630 (1988).

5. Strittmatter, W. et al. Apolipopro-tein E: high-avidity binding to beta-amyloid and increased frequency of type 4 allele in late-onset familial Alzheimer dis-ease. Proceedings of the National Academy of Sciences of theUnited States of America 90, 1977–1981 (2004).

6. Corder, E. et al. Gene dose of apolipoprotein E type 4 allele and the risk ofAlzheimer’s disease in late onset families. Science 261, 921–923 (1993).

7. Roses, A. D. Apolipoprotein E alleles as risk factors in Alzheimer’s disease. Annu. Rev. Med. 47, 387–400 (1996).

8. Bertram, L., McQueen, M. B., Mullin, K., Blacker, D. & Tanzi, R. E. Systematic meta-analyses of Alzheimer disease genetic association studies: the AlzGene database. Nat. Genet. 39, 17–23 (2007).

9. Li, Z., Shue, F., Zhao, N., Shinohara, M. & Bu, G. APOE2: protective mechanism and therapeutic implications for Alzheimer’s disease. Mol. Neurodegener. 15, 63 (2020).

10. Le Guen, Y. et al. Association of African ancestry-specific APOE missense variant R145C with risk of Alzheimer disease. JAMA 329, 551–560 (2023).

11. Farrer, L. A. et al. Effects of age, sex, and ethnicity on the association between apolipoprotein E genotype and Alzheimer disease. A meta-analysis. APOE and Alzheimer Disease Meta Analysis Consortium. JAMA 278, 1349–1356 (1997).

12. Yu, C.-E. et al. Epigenetic signature and enhancer activity of the human APOE gene. Hum. Mol. Genet. 22, 5036–5047 (2013).

13. Foraker, J. et al. The APOE gene is differentially methylated in Alzheimer’s disease. J. Alzheimers. Dis. 48, 745–755 (2015).

14. Bird, A., Taggart, M., Frommer, M., Miller, O. J. & Macleod, D. A fraction of the mouse genome that is derived from islands of nonmethylated, CpG-rich DNA. Cell 40, 91–99 (1985).

15. Gardiner-Garden, M. & Frommer, M. CpG islands in vertebrate genomes. J. Mol. Biol. 196, 261–282 (1987).

16. Saxonov, S., Berg, P. & Brutlag, D. L. A genome-wide analysis of CpG dinucleotides in the human genome distinguishes two distinct classes of promoters. Proc. Natl. Acad. Sci. U. S. A. 103, 1412– 1417 (2006).

17. Cervantes, S. et al. Genetic variation in APOE cluster region and Alzheimer’s disease risk. Neurobiol. Aging 32, 2107.e7–17 (2011).

18. Yu, C.-E. et al. Comprehensive analysis of APOE and selected proximate markers for late-onset Alzheimer’s disease: patterns of linkage disequilibrium and disease/marker association. Genomics 89, 655–665 (2007).

19. Bekris, L. M. et al. Multiple SNPs within and surrounding the apolipoprotein E gene influence cerebrospinal fluid apolipoprotein E protein levels. J. Alzheimers. Dis. 13, 255–266 (2008).

20. Cruchaga, C. et al. Cerebrospinal fluid APOE levels: an endophenotype for genetic studies for Alzheimer’s disease. Hum. Mol. Genet. 21, 4558–4571 (2012).

21. Shao, Y. et al. DNA methylation of TOMM40-APOE-APOC2 in Alzheimer’s disease. J. Hum. Genet. 63, 459–471 (2018).

22. Liu, J. et al. DNA methylation in the APOE genomic region is associated with cognitive function in African Americans. BMC Med. Genomics 11, 43 (2018).

23. Walker, R. M. et al. Identification of epigenome-wide DNA methylation differences between carriers of APOE ε4 and APOE ε2 alleles. Genome Med. 13, 1 (2021).

24. Panitch, R. et al. APOE genotype-specific methylation patterns are linked to Alzheimer disease pathology and estrogen response. Transl. Psychiatry 14, 129 (2024).

25. Mur, J. et al. DNA methylation in APOE: The relationship with Alzheimer’s and with cardiovascular health. Alzheimers Dement. (N. Y.) 6, e12026 (2020).

26. Grunau, C., Clark, S. J. & Rosenthal, A. Bisulfite genomic sequencing: systematic investigation of critical experimental parameters. Nucleic Acids Res. 29, E65–5 (2001).

27. Raizis, A. M., Schmitt, F. & Jost, J. P. A bisulfite method of 5-methylcytosine mapping that minimizes template degradation. Anal. Biochem. 226, 161–166 (1995).

28. Holmes, E. E. et al. Performance evaluation of kits for bisulfite-conversion of DNA from tissues, cell lines, FFPE tissues, aspirates, lavages, effusions, plasma, serum, and urine. PLoS One 9, e93933 (2014).

29. Sahoo, K. & Sundararajan, V. Methods in DNA methylation array dataset analysis: A review. Comput. Struct. Biotechnol. J. 23, 2304–2325 (2024).

30. Sedlazeck, F. J., Lee, H. & Darby, C. A. Schatz MC Piercing the dark matter: bioinformatics of long-range sequencing and mapping. Nat. Rev. Genet 19, 329–346 (2018).

31. Wagner, J. et al. Benchmarking challenging small variants with linked and long reads. Cell Genom. 2, 100128 (2022).

32. Lee, H. & Schatz, M. C. Genomic dark matter: the reliability of short read mapping illustrated by the genome mappability score. Bioinformatics 28, 2097–2105 (2012).

33. Sereika, M. et al. Oxford Nanopore R10.4 long-read sequencing enables the generation of near-finished bacterial genomes from pure cultures and metagenomes without short-read or reference polishing. Nat. Methods 19, 823–826 (2022).

34. Kolmogorov, M., et al. Scalable Nanopore sequencing of human genomes provides a comprehensive view of haplotype-resolved variation and methylation. bioRxivorg (2023) doi:10.1101/2023.01.12.523790.

35. Genner, R. et al. Assessing methylation detection for primary human tissue using Nanopore sequencing. bioRxiv (2024) doi:10.1101/2024.02.29.581569.

36. Billingsley, K. J., et al. Long-read sequencing of hundreds of diverse brains provides insight into the impact of structural variation on gene expression and DNA methylation. bioRxivorg (2024) doi:10.1101/2024.12.16.628723.

37. Bennett, D. A. et al. Religious Orders Study and Rush Memory and Aging Project. J. Alzheimers. Dis. 64, S161–S189 (2018).

38. Fullerton, S. M. et al. Apolipoprotein E variation at the sequence haplotype level: implications for the origin and maintenance of a major human polymorphism. Am. J. Hum. Genet. 67, 881–900 (2000).

39. Kulminski, A. M. et al. TOMM40 and APOC1 variants differentiate the impacts of the APOE ε4 allele on Alzheimer’s disease risk across sexes, ages, and ancestries. Alzheimers Dement. (Amst.) 16, e12600 (2024).

40. Kulminski, A. M. et al. Association of APOE alleles and polygenic profiles comprising APOE-TOMM40-APOC1 variants with Alzheimer’s disease neuroimaging markers. Alzheimers. Dement. 21, e14445 (2025).

41. Kulminski, A. M., Philipp, I., Shu, L. & Culminskaya, I. Definitive roles of TOMM40-APOE-APOC1 variants in the Alzheimer’s risk. Neurobiol. Aging 110, 122–131 (2022).

42. Kulminski, A. M., Philipp, I., Loika, Y., He, L. & Culminskaya, I. Haplotype architecture of the Alzheimer’s risk in the APOE region via co-skewness. Alzheimers Dement. (Amst.) 12, e12129 (2020).

43. Walton, E. et al. Correspondence of DNA methylation between blood and brain tissue and its application to schizophrenia research. Schizophr. Bull. 42, 406–414 (2016).

44. Mendonça, V., Soares-Lima, S. C. & Moreira, M. A. M. Exploring cross-tissue DNA methylation patterns: blood-brain CpGs as potential neurodegenerative disease biomarkers. *Commun*. Biol. 7, 904 (2024).

45. Taryma-Leśniak, O. et al. Methylation patterns at the adjacent CpG sites within enhancers are a part of cell identity. Epigenetics Chromatin 17, 30 (2024).

46. Vitale, D. et al. GenoTools: An open-source Python package for efficient genotype data quality control and analysis. bioRxiv (2024) doi:10.1101/2024.03.26.586362.

47. Billingsley, J. Processing Human Frontal Cortex Brain Tissue for Population-Scale Oxford Nanopore Long-Read DNA Sequencing SOP.

48. Baker, B., Abdelhalim, M. & J Billingsley, K. Processing human frontal cortex brain tissue for population-scale SQK-LSK114 Oxford Nanopore long-read DNA sequencing SOP v1. (2023) doi:10.17504/protocols.io.kxygx3zzog8j/v1.

49. Li, H. Minimap2: pairwise alignment for nucleotide sequences. Bioinformatics 34, 3094–3100 (2018).

50. Shafin, K. et al. Haplotype-aware variant calling with PEPPER-Margin-DeepVariant enables high accuracy in nanopore long-reads. Nat. Methods 18, 1322–1332 (2021).

51. Kolesnikov, A. et al. Local read haplotagging enables accurate long-read small variant calling. Nat. Commun. 15, 5907 (2024).

52. Poplin, R. et al. A universal SNP and small-indel variant caller using deep neural networks. Nat. Biotechnol. 36, 983–987 (2018).

53. Quinlan, A. R. & Hall, I. M. BEDTools: a flexible suite of utilities for comparing genomic features. Bioinformatics 26, 841–842 (2010).

54. Danecek, P. et al. Twelve years of SAMtools and BCFtools. Gigascience 10, (2021).

55. Katoh, K., Misawa, K., Kuma, K.-I. & Miyata, T. MAFFT: a novel method for rapid multiple sequence alignment based on fast Fourier transform. Nucleic Acids Res. 30, 3059–3066 (2002).

56. Price, M. N., Dehal, P. S. & Arkin, A. P. FastTree: computing large minimum evolution trees with profiles instead of a distance matrix. Mol. Biol. Evol. 26, 1641–1650 (2009).

57. Gibbs, J. R. et al. Abundant quantitative trait loci exist for DNA methylation and gene expression in human brain. PLoS Genet. 6, e1000952 (2010).

58. Patro, R., Duggal, G., Love, M. I., Irizarry, R. A. & Kingsford, C. Salmon provides fast and bias-aware quantification of transcript expression. Nat. Methods 14, 417–419 (2017).

59. Pérez-González, A. P. et al. The ROSMAP project: aging and neurodegenerative diseases through omic sciences. Front. Neuroinform. 18, 1443865 (2024).

